# Highly replicable multisite patterns of adolescent white matter maturation

**DOI:** 10.64898/2026.04.18.719321

**Authors:** Steven L. Meisler, Matthew Cieslak, Joëlle Bagautdinova, Timothy J. Hendrickson, Tanya Pandhi, Andrew A. Chen, Noah Hillman, Hamsanandini Radhakrishnan, Taylor Salo, Eric Feczko, Kimberly B. Weldon, rae McCollum, Begim Fayzullobekova, Lucille A. Moore, Lucinda Sisk, Christos Davatzikos, Hao Huang, Bárbara Avelar-Pereira, Sendy Caffarra, Kelly Chang, Philip A. Cook, Elizabeth A. Flook, Teresa Gomez, Mareike Grotheer, McKenzie P. Hagen, Zeeshan M. Huque, Iliana I. Karipidis, Arielle S. Keller, John Kruper, Audrey C. Luo, Briana Macedo, Kahini Mehta, Jamie L. Mitchell, Adam R. Pines, Laura Pritschet, Amelie Rauland, Ethan Roy, Brooke L. Sevchik, Golia Shafiei, S. Parker Singleton, Hannah L. Stone, Kevin Y. Sun, Valerie J. Sydnor, Tien T. Tong, Maya Yablonski, Jason D. Yeatman, Ariel Rokem, Russell T. Shinohara, Damien A. Fair, Theodore D. Satterthwaite

**Affiliations:** The Penn Lifespan Informatics and Neuroimaging Center, University of Pennsylvania, Philadelphia, PA, USA; Lifespan Brain Institute of Penn Medicine and CHOP, University of Pennsylvania, Philadelphia, PA, USA; Department of Psychiatry, University of Pennsylvania, Philadelphia, PA, USA; Masonic Institute for the Developing Brain, University of Minnesota Medical School, Minneapolis, MN, USA; Department of Pediatrics, University of Minnesota, Minneapolis, MN, USA; Department of Public Health Sciences, Medical University of South Carolina, SC, USA; Penn Statistics in Imaging and Visualization Endeavor, University of Pennsylvania, Philadelphia, PA, USA; Department of Biostatistics, Epidemiology and Informatics, University of Pennsylvania, Philadelphia, PA, USA; Center for AI and Data Science for Integrated Diagnostics (AI2D), Perelman School of Medicine, University of Pennsylvania, Philadelphia, PA, USA; Center for Biomedical Image Computing and Analytics, University of Pennsylvania, Philadelphia, PA, USA; Department of Radiology, University of Pennsylvania, Philadelphia, PA, USA; Department of Radiology, Children’s Hospital of Philadelphia, Philadelphia, PA, USA; Aging Research Center, Karolinska Institutet and Stockholm University, Stockholm, Sweden; Department of Biomedical, Metabolic and Neural Sciences, University of Modena and Reggio Emilia, Modena, Italy; Graduate School of Education, Stanford University, Stanford, CA, USA; Division of Developmental-Behavioral Pediatrics, Stanford University, Stanford, CA, USA; Department of Psychology, University of Washington, Seattle, WA, USA; Department of Psychology, Phillips-Universität Marburg, Marburg, Germany; Center for Mind, Brain and Behavior, Phillips-Universität Marburg, Justus-Liebig Universität Giessen, Technische Universität Darmstadt, Marburg, Germany; Lise Meitner Research Group Neuroplasticity in Development and Learning, Max Planck Institute for Human Development, Berlin, Germany; Department of Psychology and Neuroscience, Temple University, Philadelphia, PA, USA; Department of Child and Adolescent Psychiatry and Psychotherapy, University Hospital of Psychiatry Zurich, Zurich, Switzerland; Neuroscience Center Zurich, University of Zurich and ETH, Zurich, Switzerland; Department of Psychological Sciences, University of Connecticut, Storrs, CT, USA; Institute for the Brain and Cognitive Sciences, University of Connecticut, Storrs, CT, USA; Department of Neuroscience, Columbia University, New York, NY, USA; Department of Psychology, Stanford University, Stanford, CA, USA; Department of Psychiatry and Behavioral Sciences, Stanford School of Medicine, Stanford, CA, USA; Institute of Neuroscience and Medicine, Brain and Behaviour (INM-7), Research Center Jülich, Jülich, Germany; Department of Psychiatry, Psychotherapy and Psychosomatics, Faculty of Medicine, RWTH Aachen University, Aachen, Germany; Department of Psychological and Brain Sciences, University of California, Santa Barbara, CA, USA; Department of Psychiatry, University of Pittsburgh, Pittsburgh, PA, USA; eScience Institute, University of Washington, Seattle, WA, USA; Institute of Child Development, University of Minnesota, Minneapolis, MN, USA

**Author notes:** Correspondence to Steven L. Meisler (**) and Theodore D. Satterthwaite (*).

## Abstract

The Adolescent Brain Cognitive Development (ABCD) Study is the largest U.S.-based neuroimaging initiative of adolescent brain maturation. Diffusion MRI (dMRI) provides unique insights into white matter organization, yet applying advanced processing pipelines and managing technical variability across scanning environments remains challenging at scale. To address these issues, we present ABCD-BIDS Community Collection (ABCC) release 3.1.0, including a curated resource of more than 24,000 fully processed ABCD dMRI datasets. ABCC provides fully processed images, nuanced image quality metrics, advanced microstructural measures, and person-specific bundle tractography. Evaluating these rich data revealed that measures of diffusion restriction and non-Gaussianity—in particular the intracellular volume fraction from NODDI and return-to-origin probability from MAP-MRI—were highly sensitive to neurodevelopment and robust to variation in image quality. Additionally, harmonization of microstructural features markedly improved the cross-vendor generalizability of developmental effects. Together, ABCC accelerates reproducible, rigorous research on adolescent white matter development.

## Introduction

Adolescence is a critical period of brain plasticity that supports cognitive development^1–3^. However, it is also understood as a period of vulnerability to psychopathology^4,5^. As such, mapping the brain throughout adolescence provides insight into both typical and atypical trajectories of brain maturation. The Adolescent Brain Cognitive Development℠ (ABCD) Study^6^ is the largest longitudinal initiative tracking neuroimaging, behavior, cognition, and psychopathology across childhood and adolescence in the United States. Among the rich imaging data available, diffusion MRI (dMRI) is critical for characterizing the development of white matter microstructure and macrostructure^7–9^. Prior studies using ABCD dMRI data have linked white matter features to cognitive function^10,11^, the childhood environment^12,13^, and early markers of psychopathology^14–19^. Since ABCD first launched in 2015, the broader field has increasingly embraced advanced dMRI preprocessing methods^20^, more detailed quality control procedures^21^, approaches to modeling dMRI beyond the diffusion tensor, and person-specific bundle tractography. However, many of these recent methodological innovations and the extensive derivatives they produce—which may offer heightened sensitivity to white matter development^22–24^—have not been generated and publicly shared as part of the official ABCD release. Applying state-of-the-art methods across large-scale datasets like ABCD requires extensive computational resources and technical expertise, posing a significant barrier for many research groups. To accelerate rigorous and reproducible research in white matter maturation, we present the ABCD-BIDS Community Collection (ABCC)^25^ 3.1.0 data release, which includes over 24,000 ABCD dMRI datasets with analysis-ready derivatives.

This new data resource addresses four major obstacles. First, scanner-related variability may obscure generalizable developmental effects in large multisite datasets such as ABCD. Although statistical harmonization reduces mean-level scanner differences in diffusion MRI derivatives, most prior work has focused on univariate inter-site differences between regional measures^26,27^. Critically, it remains unknown whether harmonization aligns or distorts biologically meaningful spatial patterns of adolescent white matter maturation across scanning environments. White matter development follows reproducible large-scale patterns across the brain^28–30^, providing a biologically grounded benchmark for evaluating harmonization performance and microstructural model choices. However, scanner-specific factors, such as bias fields, may exhibit spatial structure that distort developmental patterns, limiting the generalizability of findings across acquisition sites. Here, we apply state-of-the-art longitudinal nonlinear harmonization and test whether reducing scanner-related variance enhances the generalizability of brain-wide developmental effects across scanning environments.

A second major challenge is that multi-shell dMRI data can be analyzed using diverse postprocessing models that yield myriad features that characterize the diffusion process. However, the relative developmental sensitivity and generalizability of these measures remain unclear. Faced with such a profusion of models and the challenges in implementing each approach, investigators typically evaluate a single or small set of measures. Historically, studies have predominantly relied on simple tensor–based measures such as fractional anisotropy (FA) and mean diffusivity (MD)^30^. More recent studies have shown that metrics from models such as neurite orientation dispersion and density imaging (NODDI)^31^, mean apparent propagator MRI (MAP-MRI)^32^, and diffusion kurtosis imaging (DKI)^33^ may be more sensitive to developmental processes than traditional tensor-derived measures^22–24^. However, these comparisons have typically been made across separate studies and cohorts, often evaluating different subsets of model features, which limits direct comparison across modeling approaches. Moreover, it remains unclear whether developmental patterns generalize across these different dMRI features or replicate consistently across scanner vendors. Here, we leverage multiple dMRI models to systematically compare the developmental sensitivity and generalizability of diverse microstructural measures in the ABCD cohort.

Third, the impact of image quality on diffusion MRI microstructural measures remains poorly characterized. These considerations are especially consequential in developmental studies such as ABCD, where dMRI image quality can vary systematically with age^34,35^ and psychopathology^36^. In longitudinal multisite consortia, age, scanning environment, and image quality may also be related. For example, earlier timepoints for a given participant—when they are younger—may be disproportionately acquired on older scanner software versions. Such alignment can complicate statistical modeling by coupling biological and acquisition-related sources of variation. Although expert manual review is often considered the gold standard for neuroimaging quality control, manually reviewing all dMRI images at the scale of ABCD is usually impractical. In recent years, numerous automated measures of dMRI image quality have been proposed, capturing features such as head motion, contrast-to-noise ratio, *q*-space consistency, and slice-wise artifacts. However, it remains unclear which of these manual or automated quality metrics capture the most microstructural variation in the context of complex multi-site longitudinal studies. We manually rated more than 3,000 ABCD dMRI images and systematically evaluated associations between dozens of measures of image quality and white matter microstructure.

Finally, we aim to lower barriers to conducting rigorous and reproducible neuroimaging research. To this end, ABCC provides fully processed, analysis-ready derivatives as both high-dimensional imaging data and tidy tabular summaries. The processing workflows used to generate these outputs—*QSIPrep*^20^ and *QSIRecon*^37^—are all open source, transparent, BIDS-compliant^38–40^, and aligned with the NMIND criteria for rigor and usability^41^. By distributing standardized derivatives generated with widely adopted community tools, ABCC reduces the need for redundant reprocessing and minimizes variability introduced by custom in-house pipelines. As such, we anticipate that the dMRI data of ABCC will accelerate robust and reproducible studies of adolescent white matter development.

## Results

We applied state-of-the-art, open-source dMRI processing pipelines, *QSIPrep* and *QSIRecon*, to longitudinal data from the ABCD study (**Fig. 1**; see **Methods**). Processing yielded 40 automated image quality metrics (IQMs; **Supplementary Table S1**), as well as many macro- and microstructural features (**Supplementary Table S2**) summarized across 67 person-specific white matter bundles (**Supplementary Fig. S1**). After quality control, 24,178 dMRI datasets across 21 sites from 10,738 participants aged 8–17 years were retained for analysis (**Table 1; Supplementary Table S3**). In addition, 33 trained reviewers manually rated 3,101 dMRI images, and a multivariate classifier was used to extend these ratings across the full cohort (**Supplementary Note S1**). These derivatives are distributed as part of ABCC release 3.1.0, with future releases incorporating additional ABCD datasets as they are acquired and processed.

**Fig. 1.**
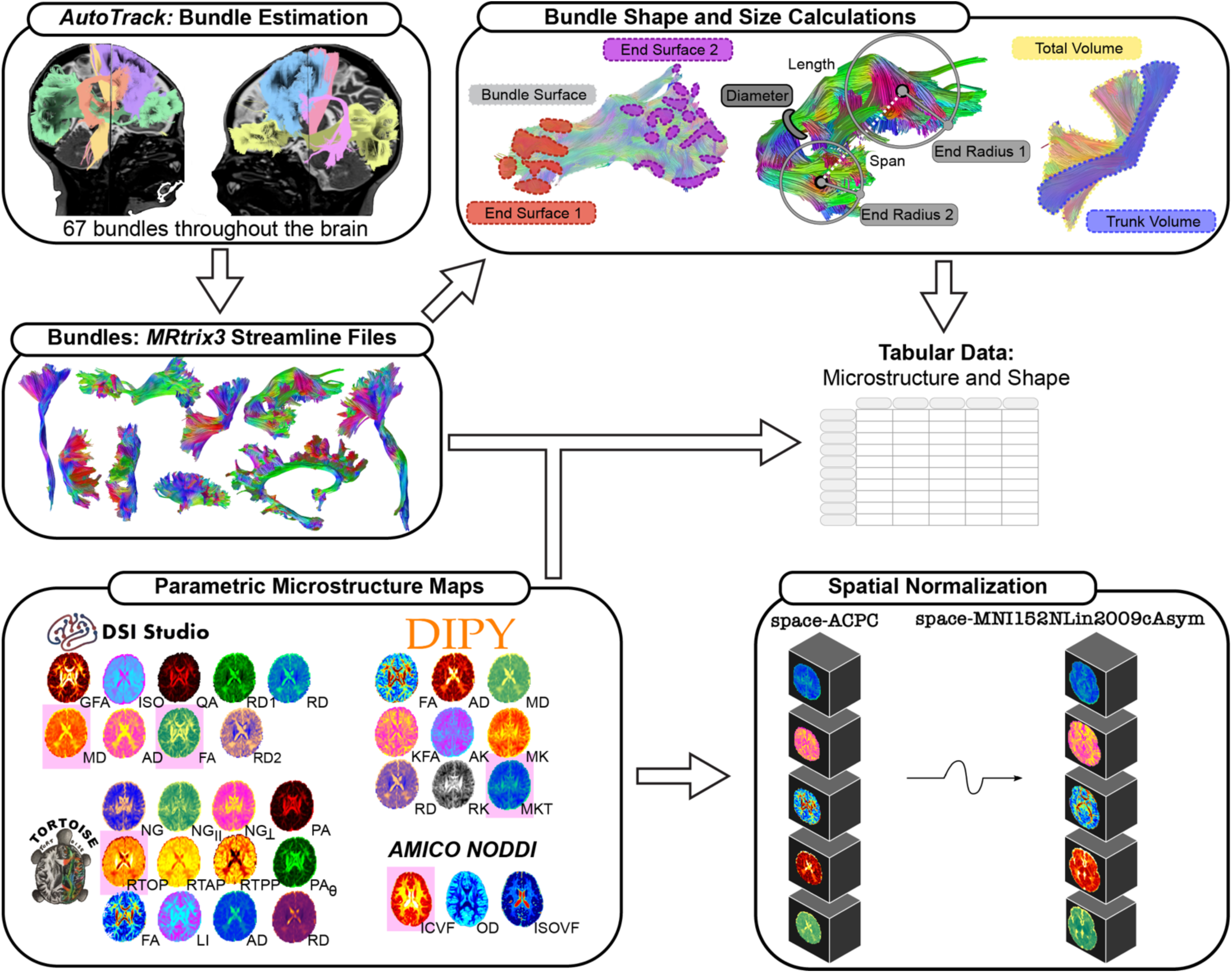
Postprocessing outputs from *QSIRecon*. *QSIRecon* was used to generate parametric microstructural maps, perform participant-specific bundle tractography, and compute bundle-wise summary statistics from ABCD diffusion MRI data. Diffusion models implemented in *DSI Studio*, *DIPY*, *TORTOISE*, and *AMICO* were applied to estimate 33 microstructural maps; the five metrics highlighted in pink are those emphasized in the primary analyses. Outputs include streamline files for 67 participant-specific white matter bundles and three-dimensional NIfTI images of each microstructural map in both participant (ACPC-aligned) and MNI152NLin2009cAsym space. Tabulated summaries of bundle-wise microstructural and macrostructural measurements are also provided. Figure adapted with permission from Cieslak et al. (2025)^42^.

**Table 1.** Summary of data. Counts of dMRI scans by ABCD study session and scanner vendor following application of inclusion and exclusion criteria. Demographic summaries include the mean age (± SD) of participants at each session and the number of male (M) and female (F) participants.

| Session | Scanner Vendor (n) |  |  | Demographics |  |  |
| --- | --- | --- | --- | --- | --- | --- |
|  | Siemens | GE | Philips | Age (Mean [SD]) | Sex (M/F) | Total |
| Baseline | 5,414 | 2,174 | 814 | 9.97 [0.63] | 4,364 / 4,038 | 8,402 |
| Year 2 | 4,335 | 1,736 | 637 | 11.99 [0.65] | 3,604 / 3,104 | 6,708 |
| Year 4 | 3,685 | 1,446 | 516 | 14.16 [0.72] | 3,024 / 2,623 | 5,647 |
| Year 6 | 2,067 | 1,043 | 311 | 16.08 [0.66] | 1,796 / 1,625 | 3,421 |
| Total | 15,501 | 6,399 | 2,278 | 12.37 [2.28] | 12,788 / 11,390 | 24,178 |

We first characterized variation in dMRI image quality across scanner vendors (Siemens, GE, and Philips) and software versions. To account for technical differences across acquisition environments, we then applied advanced nonlinear longitudinal harmonization. Using these harmonized data, we evaluated the developmental sensitivity of diffusion microstructural measures and their relationships to image quality. Throughout these analyses, we employed generalized additive mixed models (GAMMs) to account for the longitudinal data structure and flexibly model nonlinear developmental effects while minimizing overfitting. For clarity, the main results focus on 62 consistently resolved white matter bundles and five commonly used microstructural measures. Specifically, we analyzed fractional anisotropy (FA) and mean diffusivity (MD) from the *DSI Studio* tensor-fitting metric. We additionally evaluated the intracellular volume fraction (ICVF; from NODDI), return-to-origin probability (RTOP; from MAP-MRI), and mean kurtosis tensor (MKT; from DKI). These metrics span the range of diffusion modeling approaches implemented in *QSIRecon*. Results for the remaining microstructural measures are reported in the **Supplementary Materials**.

### Automated measures of image quality vary by scanner vendor and software version

Scanner heterogeneity represents a major challenge for multisite neuroimaging consortia. Even with carefully harmonized acquisition protocols, images acquired across scanner vendors (Siemens, GE, and Philips) and software versions can differ systematically in quality. *QSIPrep* produced 40 automated image quality metrics (IQMs; **Supplementary Table S1**), including neighboring dMRI correlation (NDC), dMRI contrast, contrast-to-noise ratio (CNR) within each diffusion-weighted shell, and temporal signal-to-noise ratio (tSNR) for the *b* = 0 shell. To quantify scanner-driven variation and assess the effects of preprocessing, we characterized IQMs across vendors and software versions in both raw and preprocessed data.

Substantial vendor-related variation was observed across IQMs in both raw and preprocessed data (all Kruskal–Wallis tests *p* << 0.001). Vendor effects were large for CNR across shells (**Fig. 2a**; η^2^ = 0.59–0.67), and for *b* = 0 tSNR (**Fig. 2b**; η^2^ = 0.50). Vendor differences in NDC (**Fig. 2c**; raw η^2^ = 0.39; preprocessed η^2^ = 0.68) and dMRI contrast (**Fig. 2d**; raw η^2^ = 0.17; preprocessed η^2^ = 0.15) also persisted after preprocessing. Pairwise comparisons revealed systematic patterns: Siemens scanners exhibited higher CNR, tSNR, and NDC than both GE and Philips (Wilcoxon rank-sum tests; all rank-biserial |*r*| ≥ 0.73, *p* << 0.001), indicative of higher image quality. Philips scanners showed markedly lower dMRI contrast than Siemens and GE (|*r*| = 0.73–0.80, *p* << 0.001). Differences between Siemens and GE were comparatively smaller for both raw and preprocessed dMRI contrast (both |*r*| ≤ 0.11, *p* << 0.001). As expected, preprocessing with *QSIPrep* produced consistent within-scan improvements in IQMs calculated on both raw and preprocessed data, such as NDC and dMRI contrast (**Fig. 2c,d**; paired Wilcoxon |*r*| > 0.86, *p* << 0.001).

**Fig. 2.**
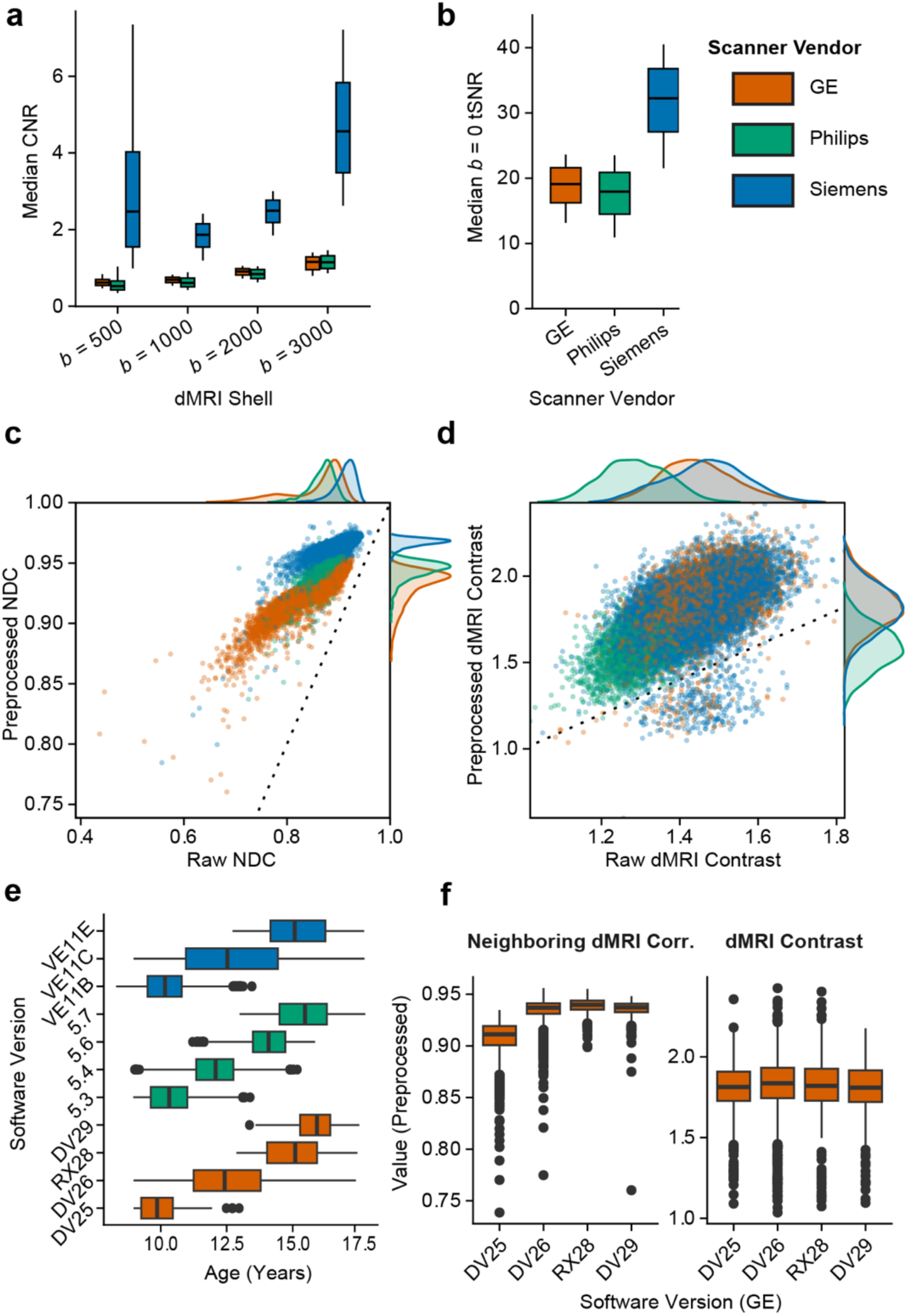
Automated image quality metrics are strongly structured by scanner vendor and software. **(a)** Shell-wise distributions of median contrast-to-noise ratio (CNR). **(b)** Median temporal signal-to-noise ratio (tSNR) in the *b* = 0 shell. **(c,d)** Neighboring dMRI correlation (NDC; left) and dMRI contrast (right) before and after preprocessing, including *B*_1_ bias correction. The dotted line denotes the identity line; values above the line indicate improved quality following preprocessing. **(e)** Distribution of ages included in each scanner software version. Boxes are colored by scanner vendor. **(f)** Distribution of preprocessed NDC (left) and dMRI contrast (right) within the GE software versions.

Scanner software versions also exhibited systematic variation. Software versions tended to vary systematically with participant age (**Fig. 2e**), reflecting scanner upgrades during the course of the study. Notably, within-vendor variation in image quality was evident across software versions (**Fig. 2f**). Within GE, for example, preprocessed NDC differed strongly across software versions (Kruskal–Wallis test; η^2^ = 0.50, *p* << 0.001), largely driven by reduced NDC in the earliest software version, DV25 (Wilcoxon rank-sum tests, all pairwise |*r*| > 0.9, *p* << 0.001). In contrast, preprocessed dMRI contrast showed substantially smaller variation across GE software versions (η^2^ = 2.82×10^-3^, *p* < 0.001). Additional IQMs and batch-stratified distributions are shown in **Supplementary Fig. S2-S9**. Together, these findings demonstrate that preprocessing substantially improves image quality, yet substantial scanner-related variation persists, underscoring acquisition heterogeneity as a fundamental challenge for large multisite diffusion MRI studies.

### Harmonization reduces mean acquisition batch effects in white matter microstructure

Although harmonization has previously been applied to ABCD diffusion MRI data, prior efforts have largely focused on cross-sectional baseline scans. Here we implemented a longitudinal nonlinear harmonization approach that preserves participant-specific effects and nonlinear developmental trajectories. To quantify acquisition-related variability, we modeled each bundle-wise microstructural metric as a function of age, sex, and acquisition batch (unique scanner device–software version combinations). Batch effect size was defined as the change in adjusted R^2^ (ΔR^2^_adj_) between the full model and a reduced model excluding the batch predictor. Analyses were performed separately for unharmonized and harmonized data.

In the unharmonized data, acquisition batch explained up to 69.8% variance in the data (**Fig. 3a**). MKT was the most susceptible to batch effects (**Fig. 3b,c**), with batch explaining 37.2% of variance on average across bundles. In contrast, RTOP was the most robust (8.2% variance explained on average). Cerebellar and subcortical fibers tended to be most susceptible to batch effects. Following harmonization, batch effects were effectively eliminated for all measures (**Fig. 3b,c**). Batch effects for all microstructural metrics are provided in **Supplementary Fig. S10.** These results demonstrate that longitudinal harmonization can be applied at scale and substantially attenuates scanner-related differences across acquisition environments.

**Figure 3.**
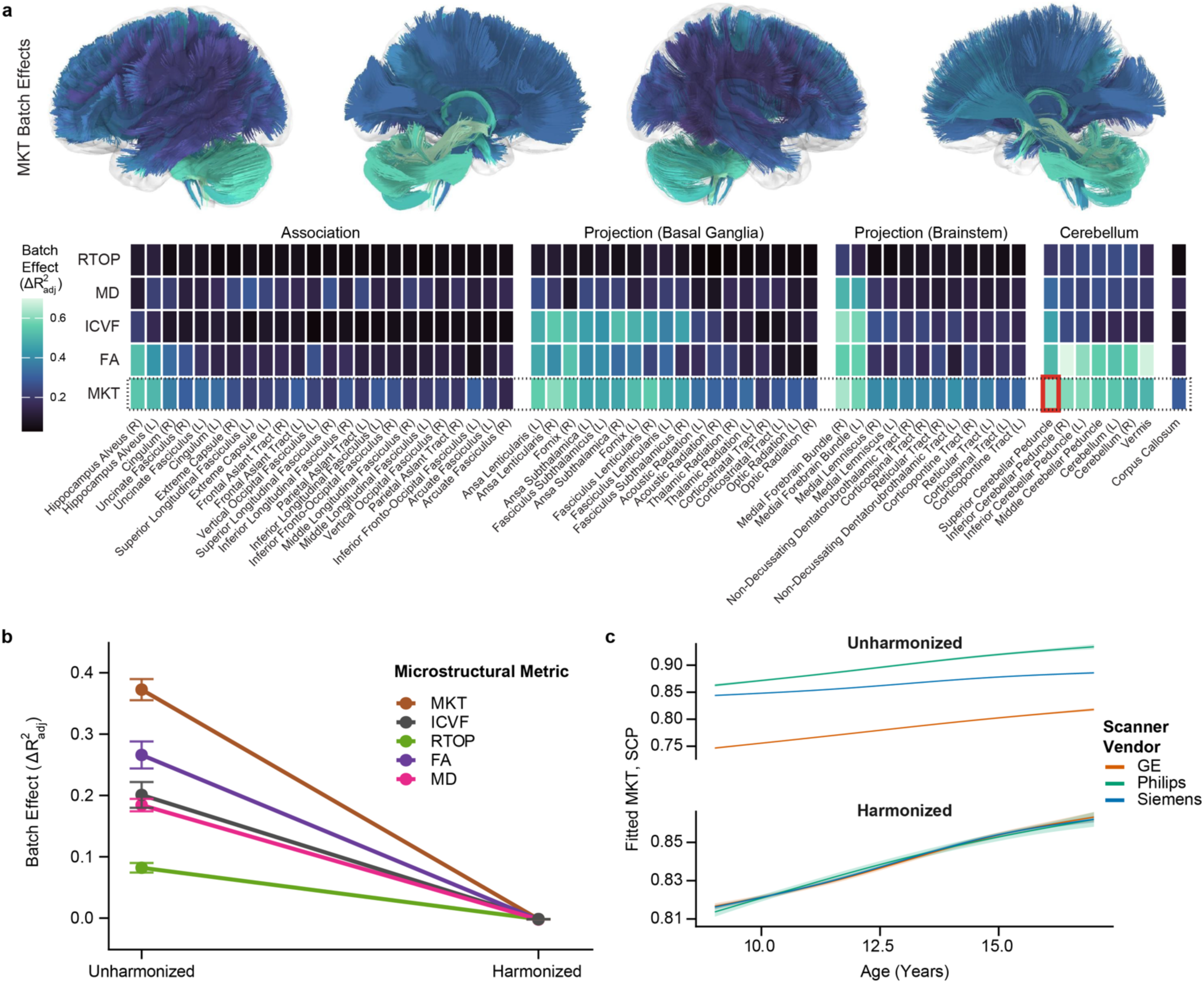
Harmonization reduces mean acquisition batch effects in white matter microstructure. **(a)** Tract plot and heatmap of batch effect sizes (ΔR^2^_adj_) in unharmonized data. Rows and columns are ordered by their mean effects, and columns are grouped by bundle category. R/L denotes right and left lateralized bundles. The red rectangle highlights MKT in the superior cerebellar peduncle (SCP). Tracts are colored by MKT batch effect size, as surrounded by a dotted rectangle in the heatmap. **(b)** Mean batch effects across bundles for each microstructural metric. Colors denote different metrics; error bars indicate the standard error of the mean. **(c)** Example developmental trajectories of MKT in the SCP before and after harmonization. Lines represent GAM-fitted mean trajectories of MKT as a function of age for each vendor; shaded regions denote 95% confidence intervals of the fitted mean.

### Advanced measures of white matter microstructure are more sensitive to development

Multi-shell dMRI enables the estimation of numerous microstructural indices beyond the diffusion tensor, yet most developmental studies have focused on a single measure or a limited subset of measures. To systematically compare their developmental sensitivity, we quantified associations between age and harmonized microstructural measures across white matter bundles. Each bundle–metric combination was modeled as a function of age and sex, and the developmental effect size was defined as the ΔR^2^_adj_ between the full model and a reduced model excluding age (**Fig. 4a**).

**Figure 4.**
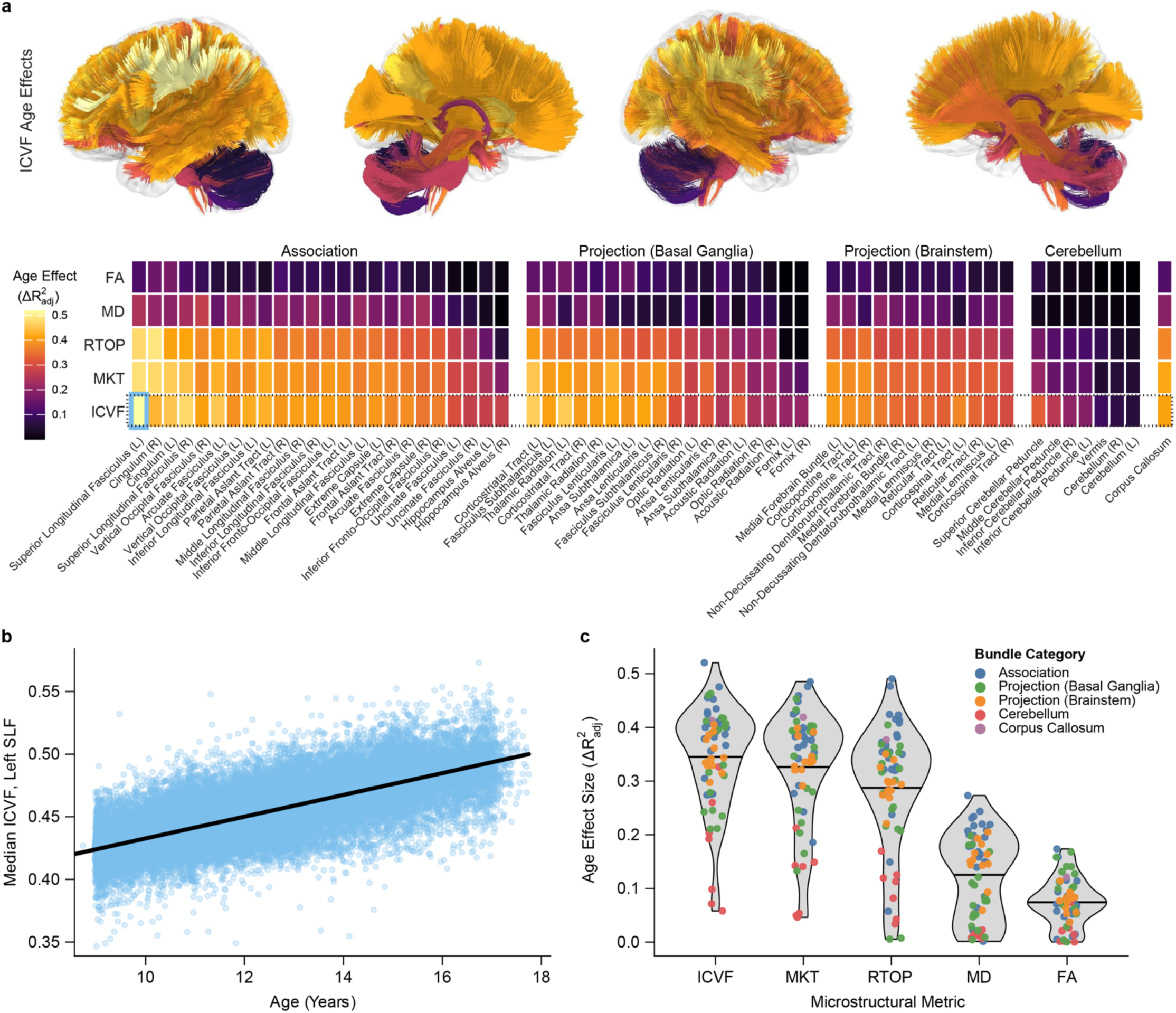
Advanced microstructural metrics are more sensitive to development. **(a)** Tract plot and heatmap of bundle-wise developmental effect sizes (ΔR^2^_adj_). Rows and columns are ordered by mean effect size, and columns are grouped by bundle category. R/L denotes right and left lateralized bundles. The blue rectangle highlights the largest observed effect (ICVF in the left superior longitudinal fasciculus; SLF). Tracts are colored by the ICVF age effect size, as surrounded by a dotted rectangle in the heatmap. **(b)** Age-related trajectory of ICVF in the left SLF from a GAM fit. The solid black curve shows the model expected ICVF value as a smooth function of age. The 95% confidence interval for the fitted mean trend is present but too narrow to be visible at this scale. **(c)** Distribution of bundle-wise age effect sizes across microstructural metrics, ordered by decreasing mean effect size across bundles. Each point represents a bundle and is colored by bundle category. Black lines indicate the mean effect across bundles for each metric.

Across microstructural metrics, the strongest developmental effects were observed for ICVF (**Fig. 4b**), followed by MKT and RTOP (**Fig. 4c**). ICVF explained 6–52% of variance across bundles (mean ΔR^2^_adj_ across bundles = 34.5%), MKT explained 5–49% (mean = 32.6%), and RTOP explained 1–49% (mean = 28.7%). In contrast, tensor-derived measures showed substantially weaker developmental sensitivity: mean diffusivity (MD) explained 0–27% of variance (mean = 12.5%), and fractional anisotropy (FA) explained 0–17% (mean = 7.4%). Developmental sensitivity also varied across bundle categories, with association bundles exhibiting the strongest age effects, and cerebellar bundles the weakest. Developmental effects for all microstructural metrics are provided in **Supplementary Fig. S11**. Together, these results emphasize that advanced multi-compartment and non-Gaussian metrics are more sensitive to development than conventional tensor-derived measures.

### Harmonization improves the generalizability of whole-brain developmental effects

Scanner effects are substantial and not uniform across the brain (**Fig. 3**). Thus, technical variation in multisite neuroimaging may distort observed patterns of development across the brain. We therefore assessed whether longitudinal harmonization improved cross-vendor generalizability of developmental effects. Within each scanner vendor, we estimated developmental effect sizes (ΔR^2^_adj_) for each bundle and microstructural metric using unharmonized and harmonized data separately. Cross-vendor correspondence was defined as the Spearman rank correlation (ρ) between vendors’ vectors of bundle-wise age effects.

Harmonization systematically increased cross-vendor correspondence for all metrics evaluated, often substantially (**Fig. 5a, Supplementary Fig. S12**). In harmonized data, all metrics showed mean harmonized correspondences greater than 0.91; MD showed the highest correspondence (mean harmonized ρ = 0.98 across vendor pairs; mean Δρ = 0.09). The most pronounced gains in generalizability following harmonization were observed for FA (mean ρ = 0.91; mean Δρ = 0.31) and MKT (mean ρ = 0.94; mean Δρ = 0.33). The largest single improvement occurred for MKT, where GE–Philips correspondence increased from 0.47 to 0.95 (Δρ = 0.48; **Fig. 5b,c**). These findings indicate that longitudinal nonlinear harmonization not only reduces mean vendor effects but also strengthens the reproducibility of spatial developmental patterns in large multisite studies.

**Figure 5.**
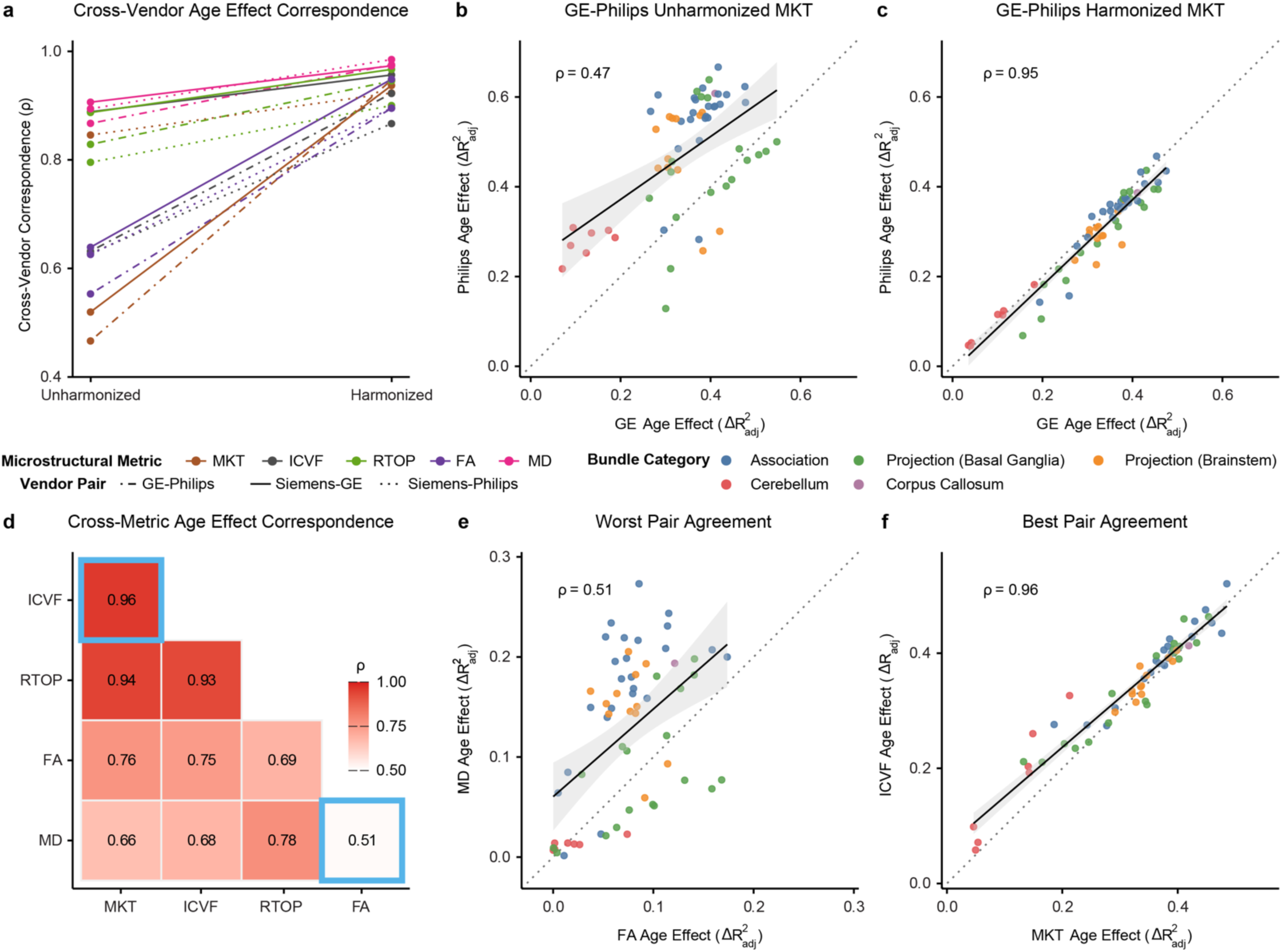
Developmental effects generalize across vendors and microstructural measures after harmonization. **(a)** Cross-vendor correspondence (Spearman ρ) of bundle-wise age effects in unharmonized and harmonized data. Colors denote microstructural metrics; line types indicate vendor pairs. **(b,c)** Illustrative plots of the largest gain in cross-vendor correspondence. Unharmonized (left) and harmonized (right) GE–Philips correspondence for MKT. Each point represents a bundle colored by bundle category. Solid lines show linear fits with 95% confidence intervals; dotted lines denote identity. **(d)** Cross-metric correspondence of developmental effects estimated from harmonized data. The strongest and weakest associations are outlined in blue. **(e,f)** Scatterplots illustrating the weakest (left) and strongest (right) cross-metric correspondences. Points represent bundles colored by category; solid lines show linear fits with 95% confidence intervals; dotted lines denote identity.

### Advanced diffusion models exhibit convergent developmental organization

Variation in the use of disparate microstructural models complicates synthesis across studies. Our results demonstrate that contemporary microstructural measures show greater developmental sensitivity than tensor-derived metrics (**Fig. 4c**). However, it remains unclear whether different models of the diffusion signal capture a shared spatial pattern of developmental effects, or whether tensor-derived effects simply reflect attenuated versions of a common developmental gradient.

We operationalized cross-metric correspondence as the Spearman rank correlation between bundle-wise age effects for each metric, estimated from the full harmonized dataset. (**Fig. 5d**; **Supplementary Fig. S13)**. Advanced measures (ICVF, RTOP, MKT) were highly correlated with one another (ρ ≥ 0.93), indicating strong agreement in the pattern of developmental effects across bundles. In contrast, tensor-derived metrics were more variable; MD showed particularly weak correspondence with other measures (mean ρ = 0.66). The strongest association was observed between ICVF and MKT (ρ = 0.96; **Fig. 5e**), whereas the weakest was between FA and MD (ρ = 0.51; **Fig. 5f**). These results indicate that developmental patterns are more consistent across advanced diffusion models than across commonly used tensor-derived measures.

### dMRI contrast captures the most microstructural variance among measures of data quality

Diffusion MRI data can be characterized using numerous automated image quality metrics (IQMs) as well as expert manual review. However, it remains unclear which measures meaningfully capture variation in data quality—or how sensitive different microstructural reconstruction methods are to variation in quality. *QSIPrep* generated 40 automated IQMs indexing diverse constructs, including head motion, signal-to-noise ratio, and *q*-space properties. Additionally, we trained 33 reviewers to rate a large subset of 3,101 images (see **Methods**). From these ratings we used a multivariate classifier to predict a quality score across the entire cohort, which was analyzed alongside the other automated IQMs.

To quantify the influence of image quality, we evaluated associations between each IQM and harmonized microstructural measures. For each bundle and microstructural metric, we fit generalized additive mixed models including age, sex, harmonization batch, and each IQM separately. Conditioning on batch allowed quality-related variance to be estimated independently of batch-driven differences. Quality effects were defined as the ΔR^2^_adj_ between the full model and a reduced model excluding the IQM predictor.

Across the five representative microstructural metrics and 62 bundles, we identified the IQM with the largest effect size for each of the 310 bundle–metric combinations (**Fig. 6a; Supplementary Fig. S14**). Preprocessed dMRI contrast (after *B*_1_ bias correction) performed the best, accounting for the largest effects in 128 combinations. In contrast, quality metrics such as mean framewise displacement and the quality classifier score rarely explained the greatest variance. Using preprocessed dMRI contrast as a representative IQM, FA was the most susceptible to quality variation (range of ΔR^2^_adj_ across bundles: −0.1–13.8%; mean ΔR^2^_adj_ = 5.3%). In contrast, ICVF was the most robust (range: −0.1–2.7%; mean = 0.8%), being on average more than sixfold less susceptible than FA (**Fig. 6b,c; Supplementary Fig. S14**).

**Figure 6.**
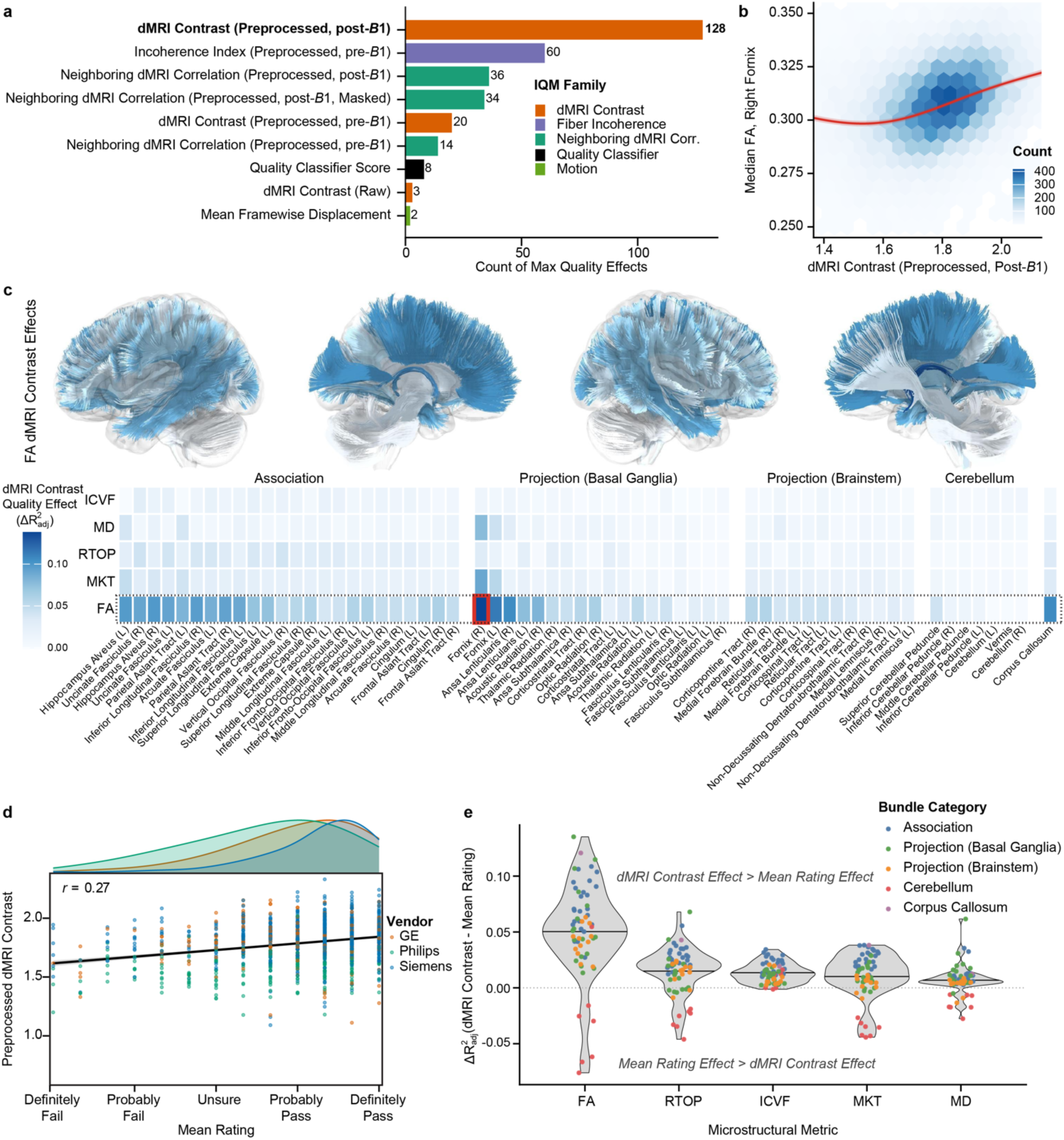
The dMRI contrast IQM captured the most variance in dMRI microstructure. **(a)** IQM winner-take-all plot. Each bar represents the number of bundle–metric combinations for which an IQM explained the greatest variance (ΔR^2^_adj_). Only IQMs with at least two instances are displayed. Bars are colored by IQM family. **(b)** Illustrative plot of preprocessed dMRI contrast against FA in the right fornix. The red curve is the GAMM fit (adjusted for age, sex, and batch), with a shaded pointwise 95% confidence band. **(c)** Tract plot and heatmap of bundle-wise quality effect sizes for preprocessed dMRI contrast across microstructural metrics. Rows are ordered by mean quality effect across IQMs; columns are ordered by mean effect size and grouped by bundle category. R/L denotes right and left lateralized bundles. The red rectangle highlights the largest observed quality effect (FA in the right fornix). Tracts are colored by the FA quality effect size, as surrounded by a dotted rectangle in the heatmap. **(d)** Scatterplot showing the relationship between the average manual rating and preprocessed dMRI contrast among the subset of images that were manually reviewed. Each color represents a different scanner vendor. **(e)** Differences in quality effect between dMRI contrast and mean manual rating among the subset of manually reviewed images. A positive value reflects a higher quality effect for dMRI contrast compared to the effect of mean manual rating. Dots are colored by bundle category. Black lines represent the mean difference in quality effects across bundles within the microstructural metric.

We further compared quality effects of preprocessed dMRI contrast and mean manual rating within the subset of manually reviewed images. dMRI contrast and manual ratings were modestly related (*r* = 0.27, *p* < 0.001; **Fig. 6d**). However, preprocessed dMRI contrast consistently showed larger quality effects than manual ratings. Overall, dMRI contrast explained more microstructural variance than manual ratings in 266 of the 310 bundle–metric combinations (**Fig. 6e**). Together, these findings indicate that dMRI contrast captures a substantial component of variability in data quality. Conversely, commonly used measures of quality such as head motion, NDC, and manual quality ratings showed markedly smaller effects. Advanced microstructural metrics were more robust to variation in image quality than fractional anisotropy, further underscoring their suitability for studies of brain development.

### Commonly used image quality metrics vary by acquisition batch and attenuate developmental effects

In developmental neuroimaging studies, image quality metrics are frequently included as covariates to mitigate spurious associations. However, age, image quality, and acquisition batch may be collinear in large multisite consortia such as ABCD. We have shown this is particularly true within GE scanners in ABCD (**Fig. 2e,f**). We therefore examined whether variation in dMRI IQMs reflected participant characteristics or instead primarily captured acquisition-related differences.

Several IQMs, including mean framewise displacement (R^2^ = 11.1%), the quality classifier score trained on manual ratings (R^2^ = 13.5%), and preprocessed NDC (R^2^ = 28.0%), showed apparent associations with age (**Fig. 7a,b**). However, variance decomposition analyses revealed that these age–IQM relationships were predominantly explained by between-batch differences rather than within-batch participant variation (≥ 82.2% of the age-IQM relationship explained by batch).

**Figure 7.**
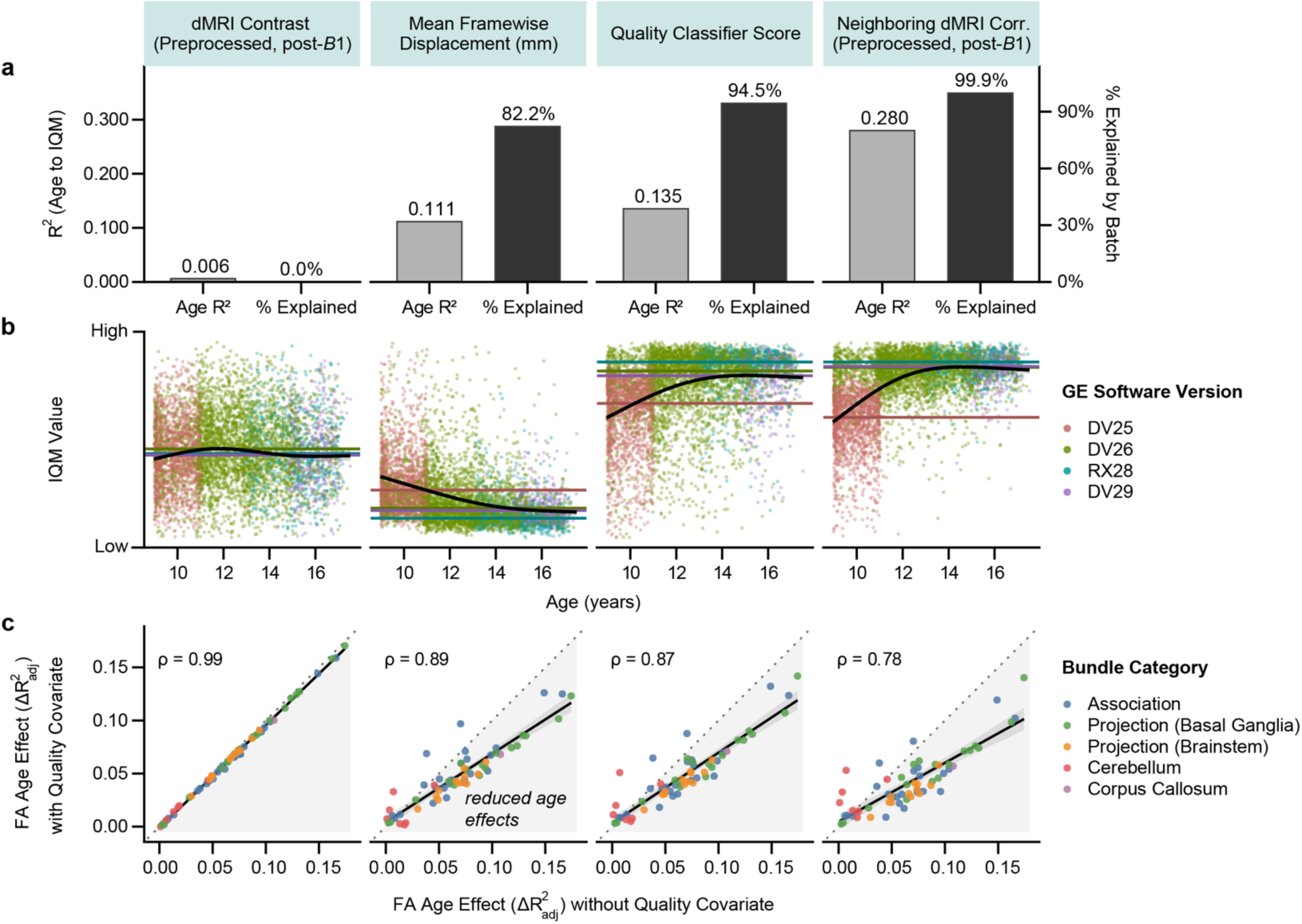
Batch-structured image quality metrics distort developmental effects in GE scanners. **(a)** Bar plots showing the age-to-IQM model R^2^ (left) and the percentage of the age–IQM relationship explained by acquisition batch (right). Percent mediation was estimated using variance decomposition comparing models with and without batch terms. **(b)** Generalized additive model (GAM) plots illustrating the relationship between age and each IQM. Each point represents a scan and is colored by scanner software version. Black curves indicate the fitted GAM with 95% confidence intervals. Horizontal colored lines indicate the median IQM value for each software version. **(c)** Comparison of bundle-wise FA age effects (ΔR^2^_adj_) estimated without (x-axis) and with (y-axis) inclusion of each IQM as a covariate. Points represent white matter bundles colored by their bundle category. The dashed line indicates the identity line (no change in estimated age effect). Gray shading highlights regions where inclusion of the IQM reduces estimated developmental effects. Spearman correlations (ρ) quantify similarity of spatial developmental patterns across bundles.

We next evaluated how including these IQMs as model covariates influenced estimates of developmental effects in white matter microstructure. Bundle-wise FA age effects were estimated in the harmonized data with and without each IQM included as a covariate. For all IQMs besides dMRI contrast, adding the metric systemically reduced age effects across bundles (**Fig. 7c**), consistent with attenuation of developmental effects due to the collinearity between age and acquisition batch. The magnitude of this attenuation varied across bundles, resulting in a loss of correspondence with developmental effects when image quality was not included as a covariate (ρ = 0.78–0.89).

Notably, preprocessed dMRI contrast showed minimal association with age or batch (age R^2^ = 0.6%, < 0.01% of the age-IQM relationship explained by batch). As such, developmental effects were nearly identical with or without a dMRI contrast covariate (ρ = 0.99). Together, these findings indicate that, in ABCD, many commonly used diffusion IQMs may reflect differences in acquisition rather than participant-level variability. In some settings, including such IQMs as model covariates can inadvertently distort biologically meaningful developmental signals.

## Discussion

Large-scale neuroimaging consortia such as ABCD provide unprecedented opportunities to characterize white matter maturation across adolescence. However, this scale also introduces important analytical challenges, including uncertainty in metric selection, scanner heterogeneity, and variation in image quality. Here, we addressed these challenges with three primary results. First, metrics from advanced diffusion models were substantially more sensitive to developmental processes than traditional tensor-derived metrics, converging on highly similar spatial patterns of maturation. Advanced measures were also more robust to image quality variation than typically used measures such as fractional anisotropy. Second, scanner-related variance was substantial and spatially structured; longitudinal harmonization not only reduced mean differences across acquisition environments but also improved the cross-vendor generalizability of developmental patterns. Finally, dMRI contrast captured the most variance in diffusion microstructure. Other commonly used quality metrics, such as head motion, contributed little explanatory value and in fact distorted estimates of developmental effects when included as model covariates. Collectively, these findings demonstrate that practical methodological choices—including the selection of microstructural metrics, harmonization strategies, and image quality covariates—can markedly impact the sensitivity and generalizability of large-scale, multisite diffusion MRI studies. To facilitate rigorous and reproducible developmental neuroimaging, we release fully processed, analysis-ready derivatives through the ABCC.

Historically, diffusion MRI studies of childhood and adolescent development have largely focused on tensor-derived metrics such as fractional anisotropy (FA) and mean diffusivity (MD), often using atlas-based analyses that obscure individual anatomical variability^30^. In contrast, we analyzed 33 harmonized microstructural metrics mapped to person-specific white matter tracts. We found that dMRI metrics from advanced models—particularly ICVF, MKT, and RTOP—were on average four- to five-fold more sensitive to developmental processes than fractional anisotropy. Importantly, these measures not only exhibited larger age-related effects but also converged on highly similar spatial patterns of maturation across white matter bundles. Although these metrics may capture distinct neurodevelopmental processes—including proliferation of neurites and increasing myelination^30,43–45^—they likely converge because they share sensitivity to restricted, non-Gaussian diffusion patterns that may not be reliably captured by tensor-based approaches. Correspondingly, tensor-derived measures showed weaker developmental effects that had different spatiotemporal patterns than the more advanced metrics. These findings highlight the value of collecting high-quality multi-shell dMRI data and fully exploiting the rich modeling framework it enables. These results also align with prior literature demonstrating heightened age sensitivity of advanced metrics among infants^42,44,46^, adolescents^23,47^, and adults^48,49^. For studies limited to diffusion tensor modeling (for example, single-shell dMRI), our results further suggest that MD—rather than FA—provides greater sensitivity to developmental change and greater robustness to variation in image quality. This observation is consistent with smaller prior studies of adolescent development^30^. However, developmental patterns estimated from tensor metrics should not be assumed to directly recapitulate those derived from advanced multi-shell diffusion models, and vice versa.

Technical variation across scanning environments can introduce substantial heterogeneity in diffusion MRI data. The utility of large multisite consortia such as ABCD therefore critically depends on the ability to harmonize data acquired across diverse scanning environments. Without effective harmonization, acquisition-related variability can dwarf more subtle biological signals of interest^26^. Here, we applied an advanced statistical harmonization approach that explicitly preserves participant-specific effects and nonlinear developmental trajectories. This approach substantially mitigated scanner-related differences in dMRI microstructural measures and increased the cross-vendor generalizability of bundle-wise developmental effects. These findings indicate that scanner-related biases are not uniformly distributed across the brain but instead exhibit spatial structure that can systematically distort developmental patterns. Accordingly, harmonization should be considered even when analyses are performed within, rather than pooled across, scanner vendors—such as when findings identified within one scanner vendor are tested for generalizability in data acquired on a different vendor. Our results extend prior work demonstrating that harmonization of ABCD dMRI data improves tractography reconstruction success and aligns bundle-wise microstructural measures across sites^27^. However, previous efforts focused on cross-sectional baseline data, excluded Philips acquisitions because of artifacts, and applied harmonization to the raw diffusion signal. In contrast, our study incorporated longitudinal data, included all scanner vendors, and applied harmonization directly to postprocessed bundle-wise features. This feature-level approach substantially reduces computational burden and facilitates application to large-scale or already-processed datasets. Together, these results underscore the importance of longitudinally informed harmonization for enabling robust multisite developmental inference in large neuroimaging consortia.

In developmental neuroimaging studies, image quality metrics are commonly included as covariates under the assumption that data quality both influences derived microstructural measures and varies systematically with participant age^50,51^. Our findings complicate this assumption in several important ways. First, metrics from advanced diffusion models showed relatively weak associations with image quality compared with fractional anisotropy, suggesting that these measures may require little or no explicit quality covariate adjustment in studies of developmental change. Second, the quality metric that most strongly tracked microstructural variation—dMRI contrast—showed minimal association with age. Individual differences in dMRI contrast likely reflect the consistency of the diffusion signal across *q*-space, arising from factors such as residual noise, large motion spikes, and scanner-related factors. In contrast, mean framewise displacement, which remains the most ubiquitous quality covariate, decreased with age but exhibited little relationship with diffusion microstructure. These findings align with and extend recent work demonstrating that modern preprocessing pipelines substantially mitigate the influence of motion on tensor-derived metrics across the lifespan^52^. Manual image assessment is often considered the gold standard for neuroimaging quality control, but it can be difficult to implement at the scale of ABCD. In our analyses, manually derived quality ratings explained relatively little variation in diffusion microstructure relative to dMRI contrast. The dMRI contrast reflects nuanced *q*-space properties of the diffusion signal that are not readily apparent through visual inspection. As a result, automated measures such as dMRI contrast may provide a principled and scalable proxy for assessing diffusion signal quality in large datasets.

Importantly, our findings suggest that in ABCD, apparent relationships between many image quality metrics and age primarily reflect acquisition-related variation rather than participant behavior. Scanner upgrades and software version changes occurred over the course of the study, resulting in partial alignment between participant age and scanning environment. Several common dMRI quality metrics showed associations with age that were predominantly explained by differences across, rather than within, acquisition batches. Notably, this pattern extended to head motion. Head motion, which is typically considered a participant-driven source of variation, was largely structured by acquisition batch in ABCD. Including these batch-structured metrics as covariates distorted estimates of developmental effects, likely due to collinearity between participant age and acquisition batch. These results suggest that routine inclusion of commonly used quality covariates may not be necessary for studies of development and, in some cases, may even distort developmental inference. This consideration is especially relevant in GE scanners—which exhibited strong collinearity among age, image quality, and acquisition batch—and for fractional anisotropy—which showed stronger associations with image quality than metrics from advanced models. dMRI contrast may capture meaningful noise-related variation in FA, but the minimal association between dMRI contrast and age suggests that including such covariates is unlikely to meaningfully alter developmental inferences.

Our study should be interpreted in the context of several limitations. First, several commonly used diffusion MRI derivatives are not currently produced as part of the ABCC pipeline. In particular, we did not include whole-brain interregional tractography, as such approaches are known to be prone to false-positive connections^53^ and impose substantial computational and storage demands at the scale of ABCD. Additionally, *QSIRecon* does not yet support restriction spectrum imaging (RSI) reconstruction^54^, precluding direct comparison with the official ABCD dMRI release. However, alternative postprocessing workflows can be applied directly to the distributed *QSIPrep*-preprocessed images, enabling future extensions of this work. Second, because optimal harmonization strategies may vary depending on the specific scientific question, we do not distribute harmonized values directly. Instead, we provide well-documented open-source code to reproduce all harmonization procedures, with the intention that these methods can be adapted to study-specific needs. Third, although longitudinal harmonization substantially reduced scanner-related variance, acquisition batch in ABCD is partially aligned with participant age due to the longitudinal study design. As a result, scanner effects and developmental effects cannot be completely disentangled, and some residual batch-related structure may remain even after harmonization.

Pre- and postprocessed dMRI data from ABCC are publicly available through release 3.1.0 to researchers with a valid Data Use Certification via the NIH Brain Development Cohorts (NBDC) Data Hub, and future releases will incorporate additional data as they are collected. Beyond the analyses presented here, our data resource provides many additional derivatives – such as measures of bundle macrostructure and volumetric microstructural maps – that support a wide range of future investigations. The end-to-end image-processing framework used here is built on widely used open-source software applicable to any BIDS-formatted dataset. As of March 2026, *QSIPrep* and *QSIRecon* have collectively been downloaded more than 58,000 times and successfully executed over 727,000 times. These tools are also the default diffusion MRI processing pipelines for the HEALthy Brain and Child Development (HBCD) study^42,55^. This broad compatibility will facilitate combining ABCD data with other cohorts using consistent processing and harmonization strategies, enabling larger and more diverse samples for studying white matter across the lifespan. Moving forward, ABCC will accelerate efforts to understand the developing brain and variation related to cognition, environment, and psychopathology.

## Methods

### Participants

A full description of the recruitment procedures, priorities, and inclusion criteria for the Adolescent Brain Cognitive Development (ABCD) Study has been previously reported^56^. Participants were initially recruited between the ages of 8 and 10 years and were invited to return for MRI assessments every two years. ABCD Release 6.0 and ABCC Release 3.1.0 include participants scanned at up to four time points, corresponding to the six-year follow-up visit at ages 15–17. All participants provided informed assent, and parents or legal guardians provided written informed consent. Study procedures at most sites were approved by a central Institutional Review Board at the University of California, San Diego; several sites obtained approval from local institutional review boards. After applying all inclusion criteria (see **Inclusion and quality control**), the final analysis sample consisted of 24,178 diffusion MRI scans from 10,738 participants (**Table 1**). Specifically, 2,892 participants contributed one scan, whereas 3,540, 3,018, and 1,288 participants contributed two, three, and four scans, respectively. The ABCC 3.1.0 release includes additional datasets that did not meet the inclusion criteria for the present analyses, as well as other modalities of data^25^. Future ABCC releases will incorporate additional ABCD data as they are acquired and processed.

### Neuroimaging acquisition and curation

Extensive details of the ABCD neuroimaging acquisition protocols have been previously described^6,57^. Data were collected across 21 sites in the United States using five scanner models from three vendors (Siemens Healthineers: Prisma and Prisma Fit; GE Healthcare: Discovery MR750 and Signa UHP; Philips: Achieva dStream and Ingenia; **Supplementary Table S3**). At each study visit, participants underwent T1-weighted (T1w) structural imaging, diffusion MRI (dMRI), and reverse phase-encoded *B_0_* field map acquisitions, among other imaging modalities. T1w images were acquired at 1 mm isotropic resolution with volumetric navigator (vNav) prospective motion correction^58^. Multi-shell dMRI scans were collected at 1.7 mm isotropic resolution with seven *b* = 0 s/mm^2^ volumes and 96 directions across four *b*-values (6 directions with *b* = 500 s/mm^2^, 15 directions with *b* = 1,000 s/mm^2^, 15 directions with *b* = 2,000 s/mm^2^, and 60 directions with *b* = 3,000 s/mm^2^). For participants scanned on Philips systems, dMRI data were acquired as two separate scans (Scan 1: 3 *b* = 0 s/mm^2^ volumes, 4 directions with *b* = 500 s/mm^2^, 7 directions with *b* = 1000 s/mm^2^, 8 directions with *b* = 2000 s/mm^2^ and 29 directions with *b* = 3000 s/mm^2^. Scan 2: 4 *b* = 0 s/mm^2^ volumes, 2 directions with *b* = 500 s/mm^2^, 8 directions with *b* = 1000 s/mm^2^, 7 directions with *b* = 2000 s/mm^2^ and 30 directions with *b* = 3000 s/mm^2^). The reverse phase-encoded *B*_0_ echo-planar imaging acquisition was obtained before the diffusion scans to enable correction of susceptibility-induced geometric distortions. Raw imaging data were curated to Brain Imaging Data Structure (BIDS) format^38,40^ and were used for both the ABCC and official ABCD data releases.

### Neuroimaging processing

For ABCC, T1w, dMRI, and fieldmap images were preprocessed with the BIDS application^39^ *QSIPrep*^20^ version 0.21.4, which is based on *Nipype* version 1.8.6^59,60^. Data were postprocessed with *QSIRecon*^37^ version 1.0.0rc2, which is based on *Nipype* version 1.9.1. These containerized software environments ensure computational reproducibility. The following paragraphs contain partially-edited text generated by *QSIPrep* and *QSIRecon*, which is distributed with a Creative Commons Zero (CC0) license with the expressed intention of being included in manuscripts.

### Anatomical preprocessing

The T1w image was corrected for intensity non-uniformity using *N4BiasFieldCorrection*^61^ from *ANTs*^62,63^ version 2.4.3, and used as an anatomical reference throughout the workflow. The anatomical reference image was reoriented into AC-PC alignment via a rigid transformation extracted from a full affine registration to the MNI152NLin2009cAsym template. A full nonlinear registration to the template from the AC-PC space was estimated via symmetric nonlinear registration (SyN) using *antsRegistration*. Brain extraction was performed on the T1w image using *SynthStrip*^64^, and automated segmentation was performed using *SynthSeg*^65,66^ from *FreeSurfer*^67^ version 7.3.1.

### dMRI preprocessing

MP-PCA denoising as implemented in *MRtrix3*’s^68^ *dwidenoise*^69^ was applied with a 5-voxel window. After MP-PCA, Gibbs unringing was performed using *TORTOISE*’s^70^ *Gibbs*^71^. Following unringing, the mean intensity of the dMRI series was adjusted so all the mean intensity of the *b* = 0 images matched across each separate dMRI scanning sequence. *FSL*’s^72^ (version 6.0.8) *eddy* was used for head motion correction and eddy current correction^73^. *eddy* was configured with a *q*-space smoothing factor of 10, a total of 5 iterations, and 1000 voxels used to estimate hyperparameters. *eddy*’s outlier replacement was run^74^. *FSL*’s *TOPUP*^75^ was used to estimate a susceptibility-induced off-resonance field based on the reverse phase-encoded *B*_0_ reference image. The *TOPUP*-estimated fieldmap was incorporated into the eddy current and head motion correction interpolation. Final interpolation was performed using the Jacobian modulation method. The dMRI time series was rigidly aligned to AC-PC orientation to match the T1w image, preserving the original 1.7 mm isotropic voxel resolution. *B*_1_ field inhomogeneity was then corrected using *dwibiascorrect* from *MRtrix3* with the N4^61^ algorithm.

### Automated image quality measures

A set of 40 automated image quality metrics (IQMs) were generated by *QSIPrep* (**Supplementary Table S1**), as detailed below. Vendor-wise differences in IQMs were tested with Kruskal–Wallis tests and post hoc pairwise Wilcoxon rank sum tests. A quality classifier score was also calculated after training a multivariate model to predict expert manual ratings based on automated IQMs (see **Supplementary Note S1**). Pre-to-post differences in IQMs were tested with paired Wilcoxon signed-rank tests. Correlations between IQMs are reported in **Supplementary Fig. S15**.

### Motion and registration

Seven IQMs were derived to index in-scanner motion and registration quality. The average and maximum framewise displacement values were recorded, following the definition described by Power and colleagues (2012)^76^. Additionally, the largest values of translation and rotation were recorded, along with the maxima of their derivatives. To gauge T1w-to-dMRI co-registration quality, the difference in spatial overlap between the anatomical brain mask and the dMRI brain mask was calculated using the Dice coefficient.

### Contrast-to-noise ratio

The average, median, and standard deviation of the voxel-wise contrast-to-noise ratio was calculated within each of the four dMRI diffusion-weighted shells (i.e., excluding *b* = 0) after eddy current and motion correction, totaling 12 IQMs. Additionally, the average, median and standard deviation of the voxel-wise temporal signal to noise ratio (tSNR) of the *b* = 0 volumes was calculated. The voxel-wise values are calculated as part of *FSL*’s *eddy* command within *QSIPrep*.

### Fiber coherence

The fiber coherence and incoherence indices, as described in Shilling and colleagues (2019)^77^ were calculated to measure fixel-by-fixel connectivity and gauge the validity of the gradient table in the raw data and in the preprocessed dMRI after accounting for transformations from motion correction and AC-PC alignment^78^. These two indices were derived from the raw signal, as well as preprocessed image before and after *B*_1_ bias correction, for a total of 6 IQMs.

### dMRI signal similarity and dissimilarity

A total of 12 IQMs were generated to quantify expected dMRI signal similarity between adjacent slices and gradient directions, as well as expected dMRI signal dissimilarity between perpendicular gradient directions. First, the number of poor-quality slices was calculated for the raw data, as well as the preprocessed dMRI before and after *B*_1_ bias correction for a total of three IQMs. A slice was considered low quality if the sum of its voxel-wise intensity differences from the immediately preceding and following slices exceeded twice the voxel-wise intensity difference between those two neighboring slices. Finally, the average neighboring dMRI correlations (NDC)^79^ and dMRI contrasts were calculated. NDC for a given volume is quantified as the voxel-wise correlation between that volume and its nearest neighboring volume in *q*-space. A value closer to 1 indicates a higher degree of expected signal similarity, denoting higher quality. dMRI contrast for a volume is quantified as NDC divided by the voxel-wise correlation between that volume and the volume with the most perpendicular direction in *q*-space. A higher dMRI contrast value indicates a higher degree of expected signal dissimilarity, denoting higher quality. These indices were derived on the raw and preprocessed dMRI series before and after *B*_1_ bias correction. Additionally, NDC was calculated on both the entire 3D volume space and restricted to the brain mask; in total this yielded 9 IQMs.

### dMRI postprocessing

Several dMRI reconstruction models were run as part of *QSIRecon*, resulting in a broad set of derived microstructural maps (**Fig. 1, Supplementary Table S2**). All maps described below were produced in participant-specific dMRI space (AC-PC orientation), as well as in MNI152NLin2009cAsym space using nearest neighbor interpolation to facilitate mass univariate analyses. Nearest-neighbor interpolation was used when warping scalar maps to standard space to avoid spatial smoothing artifacts (e.g., ringing in the corpus callosum) introduced by higher-order interpolation during nonlinear transformations. 33 of these metric maps were evaluated in this study, and 5 are focused on in the main text. Postprocessing software bundled in *QSIRecon* included *DIPY* version 1.8.0, *TORTOISE* version 4 (5d0c688e), *DSI Studio* version “Hou” (03476a67), and *AMICO* version 2.0.3.

### Diffusion tensor imaging and diffusion kurtosis imaging

The diffusion tensor imaging (DTI) model characterizes the 3D Gaussian diffusion process in each voxel by estimating a second-order tensor from the diffusion-weighted signal. The diffusion kurtosis imaging (DKI) model extends the tensor formulation by quantifying the degree to which water diffusion deviates from Gaussian behavior. This is achieved by additionally modeling a fourth-order kurtosis tensor that captures microstructural complexity not represented by DTI. Estimates of DTI parameters from a DKI model may be more reliable than from a standalone DTI model^80^.

In ABCC, the DTI model was fitted using three different software. The inner-shell tensor fit (*b* ≤ 1000 s/mm^2^) was computed twice. *DSI Studio* used an ordinary least squares approach, while *TORTOISE* used a weighted linear least squares^81^ approach, in which the tensor parameter estimation is weighted proportional to the estimated CNR. Additionally, a DKI fit was run with *DIPY* using the full dMRI data, producing both DKI and DTI derivatives. The parametric microstructural maps output by each software varied (**Fig. 1, Supplementary Table S2**), but the total set of DTI maps we analyzed included fractional anisotropy (FA), mean diffusivity (MD), axial diffusivity (AD), average radial diffusivity (RD), individual radial diffusivity components (RD1 and RD2), and lattice index (LI). DKI maps included kurtosis fractional anisotropy (KFA), mean kurtosis (MK), mean kurtosis tensor (MKT), axial kurtosis (AK), and radial kurtosis (RK). Other derived DTI maps, such as the average *b =* 0 amplitude (AM) and individual tensor components (e.g., *T_XX_*) are included in the data release but are not considered microstructural indices. These auxiliary metrics were not considered in the present analyses. A comparison of FA and MD results across the three tensor-fitting workflows is provided in **Supplementary Fig. S16**).

### Generalized *q*-sampling imaging

Generalized *q*–sampling imaging (GQI) is a model-free reconstruction approach that estimates the diffusion orientation distribution function (ODF) directly from the *q*-space signal without assuming Gaussian diffusion or fitting a tensor model^82^. By leveraging the Fourier relationship between the diffusion signal and the spin displacement distribution, GQI recovers high-angular resolution ODFs that capture complex fiber configurations—including crossing, fanning, and branching—that are not resolvable with traditional tensor-based methods. These ODFs were used to derive quantitative indices such as quantitative anisotropy (QA), generalized fractional anisotropy (GFA), and isotropic diffusion (ISO). The helix angle (HA) was also calculated, but is not analyzed in this study. In *QSIRecon*, GQI modeling is performed by *DSI Studio*.

### Mean apparent propagator MRI

Mean apparent propagator MRI (MAP-MRI)^32^ models the full three-dimensional probability distribution of water molecule displacements, named the ensemble average propagator (EAP). MAP-MRI does not impose Gaussian assumptions and instead captures parametric features of the propagator, enabling sensitive characterization of microstructural complexity, directional dependence, and restrictions to diffusion. The method yields a suite of microstructural maps derived from the EAP that reflect different aspects of tissue organization and diffusion. MAP-MRI was run with *TORTOISE* and produced microstructural maps including non-Gaussianity (NG), its directional components parallel and perpendicular to the principal diffusion axis (NG∥ and NG⊥), propagator anisotropy (PA), the angular distance between the estimated EAP and an isotropic EAP (PAθ), as well as maps of return-to-origin (RTOP), return-to-axis (RTAP), and return-to-plane (RTPP) probabilities.

### Neurite orientation dispersion and density imaging

Neurite Orientation Dispersion and Density Imaging (NODDI)^31^ is a biophysical model that characterizes microstructural features of brain tissue by estimating the relative contributions of intra-neurite, extra-neurite, and isotropic (CSF-like) compartments. The model provides indices such as intracellular volume fraction (ICVF, synonymous with neurite density index or NDI), orientation dispersion index (ODI), and isotropic volume fraction (ISO), which reflect neurite density, angular dispersion, and free-water content, respectively. We used the Accelerated Microstructure Imaging via Convex Optimization (*AMICO*)^83^ framework to fit NODDI with fixed parallel intrinsic and isotropic diffusivities of 0.0011 and 0.003 mm²/s, respectively^84^.

### Tractography and tractometry

A set of 67 white matter bundles across the brain were segmented for each dMRI image using *DSI Studio*’s *AutoTrack*^85^ (**Supplementary Fig. S1**). These bundles span several major anatomical categories, including association, projection, commissural, cerebellar, and brainstem pathways. This pipeline uses quantitative anisotropy and isotropic diffusion maps (both derived from GQI) to inform a non-linear registration from MNI152NLin2009aAsym space to participant-space. Bundle priors were based on a population-average white matter atlas^86^. For each bundle, deterministic tractography seeds were placed around the atlas-defined bundle region mapped to subject-space and propagated until they terminated. Streamline propagation was based on fiber orientation distribution functions (FODFs) derived from multi-shell multi-tissue constrained spherical deconvolution (MSMT-CSD)^87^, as implemented in *MRtrix3*. The spherical harmonics from the *MRtrix3* FODFs were resampled to the default direction set used by *DSI Studio* for cross-software compatibility. Bundle-wise averages and medians for every metric map produced by *QSIRecon* were calculated. For this study, we focused on the bundle-wise median values. Additionally, bundle-wise macrostructural shape metrics^85^, such as volume and surface area, were calculated. These bundle-wise summary measures were saved as tabular data.

### Inclusion and quality control

Following previously described procedures^57^, a subset of ABCD images were manually reviewed by the ABCD Data Analysis, Informatics & Resource Center. These initial quality assurance checks confirmed that each dMRI scan was complete and had an accompanying *B_0_* map for susceptibility distortion estimation. Scans were also flagged if derived dMRI maps (e.g., mean *b = 0* image, FA, MD) exhibited artifacts. Any failing images were excluded before curation and processing and were not included in either the official ABCD or ABCC data releases.

ABCC release 3.1.0 includes 24,774 datasets from 10,844 participants with both *QSIPrep* and *QSIRecon* outputs on the NIH Brain Development Cohorts (NBDC) Data Hub. For this study, we applied additional quality assurance protocols. A set of five bundles, including the bilateral dentatorubrothalamic tracts, bilateral corticobulbar tracts, and the anterior commissure, had relatively high failure rates of between 257 and 1,055 instances each. We did not analyze these five bundles. Furthermore, datasets were excluded for any of the following reasons: (1) They were recommended for exclusion by the ABCD fast track quality control protocol (160 datasets excluded); (2) they had missing or invalid scanner data (50 datasets excluded); (3) reconstruction failed for any of the remaining 62 bundles (300 datasets excluded); (4) they exhibited implausible tract geometry statistics (see **Supplementary Note S2**; 310 datasets flagged for exclusion; 77 non-overlapping datasets with previous exclusion criteria); or (5) they belonged to a harmonization batch containing fewer than 15 images (see below; 9 datasets excluded). Our final sample consisted of 24,178 datasets from 10,738 participants with complete imaging and phenotypic data.

### Manual quality rating

We trained reviewers to manually rate the quality of a subset of 3,101 ABCC preprocessed dMRI images. We used these manual ratings to train a classifier to predict quality ratings across the entire dataset. To ensure consistency, all raters completed standardized training. This involved engaging with reading materials and demonstration videos highlighting common artifacts. Reviewers also completed a pre-rating quiz, in which they were asked to all rate the same 10 images. Although there is no ground truth, an answer key (written by authors S.L.M. and M.C.) provided explanations for how expert reviewers would likely rate the images in the quiz set. The training materials may be found at https://osf.io/89uya/overview?view_only=2559ba4e71b54c07bff70aef302540d5. Each reviewer assigned every image an integer score from –2 (“definitely fail”) to +2 (“definitely pass”), with 0 indicating uncertainty. All assessments were made using the browser-based *dmriprep-viewer* interface^21^, which presents the full dMRI time series alongside framewise displacement and head-translation plots, slice-wise noise timeseries, registration overlays, and color-coded fractional anisotropy maps. In general, reviewers were instructed to focus on overall *practical usability* of the dMRI. In other words, a single noisy volume might be acceptable if the remainder of the series and fractional anisotropy map were clean. However, issues that were pervasive across the entire timeseries, such as poor brain masking, severe persistent motion corruption, or systemic noise, warranted a failure. While reviewers could assess visual depictions of motion via the interface, they were blinded to all other quantitative IQMs.

A total of 33 reviewers were employed to rate images. First, a team of 23 trained reviewers evaluated images acquired on Siemens scanners. During this stage 1,771 Siemens dMRI images from the baseline session were randomly selected so that, across all combinations of the 23 reviewers, each image received exactly three independent ratings. Each reviewer rated 231 images equally distributed across 7 quality percentile bins (indexed by the unmasked processed NDC before *B*_1_ bias correction), and each reviewer was paired with each other reviewer 21 times to gauge inter-rater reliability. In a separate stage, 21 raters (11 of these raters overlapped with the Siemens reviewers) reviewed a total of 1,330 dMRI images, with 665 images coming from each GE and Philips across all timepoints. Each reviewer rated 190 images, with the GE-to-Philips split being roughly half, again divided across 7 quality percentile bins. Each image was rated by 3 reviewers, and raters were paired with one another 19 times to gauge inter-rater reliability. Inter-rater disagreement for a particular image and pair of reviewers was quantified as the absolute difference in rating scores (**Supplementary Fig. S17**). Of the 3,101 scans, 2,829 were included in the total cohort analyzed in this study. However, all 3,101 scans were used to train the quality classifier (see **Supplementary Note S1**).

### White matter feature harmonization

To correct for technical variation across scanning environments, we applied a harmonization procedure to the bundle-wise median values for each derived dMRI microstructural metric. We used longitudinal *ComBat-GAMM*, an extension of *longComBat*^88^, as implemented in the R package *ComBatFamily* (https://github.com/Nhillman19/ComBatFamily commit f8f9f3d, based off of https://github.com/andy1764/ComBatFamily version 0.2.1). We defined a harmonization batch as each unique scanner serial number and major software version combination with at least 15 images. This resulted in 60 unique batches (**Supplementary Table S3**). During harmonization, we used generalized additive mixed models (GAMM) to remove the mean effect of batch while preserving the fixed effects of age and sex, including participant-specific random intercepts and age slopes to account for the longitudinal data structure. The age fixed effect was modeled as a penalized spline (maximum basis dimension *k* = 4). Harmonization for each microstructural metric was performed jointly across all bundles to take advantage of the empirical Bayesian estimation of average batch differences.

### Quantifying the effect of batch on dMRI microstructure

We quantified the variance attributable to harmonization batch before and after harmonization. For each bundle and microstructural metric, we fit generalized additive mixed models (GAMMs) relating the median bundle value to age, sex, and acquisition batch, with person-specific random intercepts and age slopes to account for the longitudinal data structure. Age was modeled as a penalized spline (maximum basis dimension *k* = 4). The batch effect size for each bundle–metric combination was defined as the change in adjusted R^2^ (ΔR^2^_adj_) between the GAM component of the full model and a reduced model excluding the batch term. This procedure was performed separately for unharmonized and harmonized data.

### Developmental sensitivity of dMRI microstructure

We quantified developmental sensitivity of diffusion microstructural measures during childhood and adolescence. For each bundle and microstructural metric, we fit GAMMs relating the median bundle value to age while controlling for sex, with person-specific random intercepts and age slopes to account for the longitudinal data structure. Age was modeled as a penalized spline (maximum basis dimension *k* = 4). The age effect size for each bundle–metric combination was defined as the ΔR^2^_adj_ between the GAM component of the full model and a reduced model excluding the age term. Additional models were also fit including individual image quality metrics (IQMs) as covariates to evaluate how controlling for image quality influenced estimates of developmental effects.

### Cross-vendor and cross-metric correspondence of developmental effects

We tested whether harmonization improved the spatial correspondence of developmental effects between scanner vendors. Within each scanner vendor, we fit GAMMs relating median bundle values to age and sex, with random effects for participants. For Philips scanners, too few participants (549 out of 1,259) had longitudinal data to reliably estimate random slopes, so models included only person-specific random intercepts. For Siemens and GE scanners, both random intercepts and age slopes were included. Age was modeled as a penalized spline (maximum basis dimension *k* = 4). For each vendor, microstructural metric, and bundle, we calculated the developmental age effect size as the ΔR^2^_adj_ between the GAM component of the full model and a reduced model excluding the age term. For each metric and pair of vendors (Siemens–Philips, Siemens–GE, and GE–Philips), we then calculated Spearman rank correlations between the vendors’ vectors of bundle-wise age effect sizes. This procedure was performed separately on unharmonized and harmonized data.

We also characterized the correspondence of developmental patterns across microstructural metrics. Using the age effect sizes estimated from the harmonized data pooled across vendors, we calculated Spearman rank correlations between each pair of metrics’ vectors of bundle-wise age effect sizes.

### Sensitivity of dMRI microstructure to image quality

We evaluated the susceptibility of derived diffusion microstructural metrics to variation in image quality. For each bundle and microstructural metric, we fit GAMMs relating the harmonized median bundle value to each image quality metric (IQM), while controlling for age, sex, and harmonization batch, with person-specific random intercepts and age slopes to account for the longitudinal data structure. Conditioning on harmonization batch ensured that quality-related variance was estimated independently of acquisition-related differences across scanners and software versions. The effects of age and the IQM were modeled as penalized splines (maximum basis dimension *k* = 4). Our analysis space included 33 *QSIRecon*-derived microstructural metrics as dependent variables and 41 image quality metrics as predictors, comprising the 40 automated IQMs generated by *QSIPrep* and the quality classifier score derived from manual ratings. For each model, we additionally fit a reduced model excluding the IQM term. The quality effect size for each bundle–metric combination was defined as the ΔR^2^_adj_ between the GAM components of the full and reduced models.

For the subset of 2,829 manually rated images, we additionally calculated quality effects for the mean manual rating and preprocessed dMRI contrast. Because the manually rated subset contained limited longitudinal observations, these analyses were performed using generalized additive models without random effects.

### Variance decomposition of age–image quality relationships

To determine whether apparent age–IQM associations reflected biological developmental effects or instead arose from alignment between participant age and acquisition batch, we performed variance-decomposition analyses within a generalized additive modeling framework. For each IQM, we fit three models: (i) an age-only model *(IQM ∼ s(age)*), (ii) a batch-only model (*IQM ∼ batch*), and (iii) a combined model (*IQM ∼ s(age) + batch*). We quantified the proportion of age-related variance explained by batch by comparing the variance explained in the age-only model with the age-related variance retained after conditioning on batch (R^2^_full_ − R^2^_batch_). Finally, to assess the downstream impact of quality adjustment on developmental inference, we calculated Spearman rank correlations between bundle-wise age effect sizes estimated from models with versus without added image quality covariates.

## Data availability

The present work used raw data from the Adolescent Brain Cognitive Development (ABCD) Study through Release 6.0 (https://doi.org/10.82525/jy7n-g441). Raw ABCD data and the ABCC processed derivatives described in this article are available through the NIH Brain Development Cohorts Data Hub (NBDC; https://www.nbdc-datahub.org/) to researchers who obtain and adhere to a Data Use Certification.

## Code availability

Container images for *QSIPrep* and *QSIRecon* are available on DockerHub (https://hub.docker.com/r/pennlinc/qsiprep/tags and https://hub.docker.com/r/pennlinc/qsirecon/tags), and can be executed using Docker^89^ or Singularity/Apptainer^90^. Study-specific analysis code and instructions for reproducing the analyses presented in this study are available at https://github.com/PennLINC/Meisler_ABCD_dMRI. Manual rating training material may be found at https://osf.io/89uya/overview?view_only=2559ba4e71b54c07bff70aef302540d5.

## Acknowledgements

This research was supported by the National Institutes of Health (T32MH019112 to S.L.M.; R37MH125829 to D.A.F. and T.D.S.; 2R01MH112847 to R.T.S. and T.D.S.; 2R01MH113550 to T.D.S.; R01MH123550 to R.T.S.; F30MH138048 to K.Y.S.; RF1MH121868, RF1MH121867, RF1MH126699, R01AG060942, U19AG066567, R01EY033628, and R01EB027585 to A.R.; T32MH016804 and T32MH018951 to V.J.S.; R01EB031284 to H.H.; F31MH136685 to J.B.; R01DA056499 and R25DA061823 to E.F.). S.L.M. was additionally supported by the Hartwell Foundation. G.S. was supported by a postdoctoral fellowship from the Canadian Institutes of Health Research (CIHR). A.S.K. was supported by a NARSAD Young Investigator Award from the Brain & Behavior Research Foundation. S.C. was supported by the European Union’s Horizon 2020 research and innovation program under the Marie Skłodowska-Curie grant agreement No. 837228 and PRIN PNRR P2022SMEJW. J.K. (DGE-2140004) and Z.M.H. (2038235) were supported by the National Science Foundation Graduate Research Fellowship Program. L.M.S. was supported by a National Science Foundation SBE Postdoctoral Research Fellowship (2507497). A.R. (269953372/GRK2150) and M.G. (EXC 3066/1 “The Adaptive Mind”, Project No. 533717223) were supported by the German Research Foundation (Deutsche Forschungsgemeinschaft, DFG). M.G. is further supported by the Lise Meitner Excellence Program of the Max Planck Society and the European Union (ERC, WRAPPED, 101161197). Views and opinions expressed are those of the authors only and do not necessarily reflect those of the European Union or the European Research Council. Neither the European Union nor the granting authority can be held responsible for them.

Data used in the preparation of this article were obtained from the *Adolescent Brain Cognitive Development*℠ (*ABCD*) Study, held in the NIH Brain Development Cohorts Data Sharing Platform. This is a multisite, longitudinal study designed to recruit more than 10,000 children age 9–10 and follow them over 10 years into early adulthood. The ABCD dataset grows and changes over time. The raw ABCD data used in this report came from ABCD release 6.0: https://doi.org/10.82525/jy7n-g441.

The ABCD Study® is supported by the National Institutes of Health and additional federal partners under award numbers U01DA041048, U01DA050989, U01DA051016, U01DA041022, U01DA051018, U01DA051037, U01DA050987, U01DA041174, U01DA041106, U01DA041117, U01DA041028, U01DA041134, U01DA050988, U01DA051039, U01DA041156, U01DA041025, U01DA041120, U01DA051038, U01DA041148, U01DA041093, U01DA041089, U24DA041123, U24DA041147. A full list of supporters is available at https://abcdstudy.org/about/federal-partners/.

## Contributions

S.L.M., M.C., and T.D.S. planned and designed the analyses; E.F., K.B.W., T.J.H., r.M., B.F., T.P., L.A.M., and D.A.F. curated, stored, and published the data; S.L.M. and M.C. processed the neuroimaging data with software developed by S.L.M., M.C., and T.S.; S.L.M. ran statistical analyses; S.L.M. and J.B. generated figures; S.L.M. and M.C. prepared training materials for manual data quality raters. A.A.C, N.H., and R.T.S guided usage of harmonization software. C.D. provided computing resources. S.L.M., M.C., H.R., B.A.-P., J.B., S.C., K.C., P.A.C., E.A.F., T.G., M.G., M.P.H., Z.M.H., I.I.K., A.S.K., J.K., A.C.L., B.M., K.M., J.L.M., A.R.P., L.P., A.R., E.A.R., B.S., G.S., P.S., H.L.S., K.Y.S., V.J.S., T.T., M.Y., and A.R. were manual data quality raters; S.L.M. wrote the original draft and all authors reviewed and revised the final draft.

