## Supplemental Materials for "Highly replicable multisite patterns of adolescent white matter maturation"

### Supplementary Materials for Meisler et al., ABCC dMRI Manuscript

| Image Quality Metric (IQM) | *QSIPrep* Variable Name | Higher Quality Indication |
|---|---|---|
| **Head motion metrics** | | |
| Mean Framewise Displacement | `mean_fd` | Lower value (less head motion) |
| Maximum Framewise Displacement | `max_fd` | Lower value (less extreme motion) |
| Maximum Translation | `max_translation` | Lower value (less motion) |
| Maximum Rotation | `max_rotation` | Lower value (less rotation) |
| Maximum Relative Translation | `max_rel_translation` | Lower value (less motion variability) |
| Maximum Relative Rotation | `max_rel_rotation` | Lower value (less rotation variability) |
| **Registration quality** | | |
| T1–dMRI Dice Distance | `t1_dice_distance` | Lower value (better spatial overlap) |
| **Signal- and contrast-to-noise metrics** | | |
| Mean tSNR ($b$ = 0) | `CNR0_mean` | Higher value (better contrast) |
| Mean CNR (Shell 1–4) | `CNR[1-4]_mean` | Higher value |
| Median tSNR ($b$ = 0) | `CNR0_median` | Higher value |
| Median CNR (Shell 1–4) | `CNR[1-4]_median` | Higher value |
| **Signal stability metrics** | | |
| tSNR SD ($b$ = 0) | `CNR0_standard_deviation` | Lower value (more consistent contrast) |
| CNR SD (Shell 1–4) | `CNR[1-4]_standard_deviation` | Lower value |
| ***q*-Space similarity metrics** | | |
| Neighboring dMRI Correlation | `[raw|t1|t1post]_neighbor_corr` | Higher value, closer to 1 (higher neighboring dMRI similarity) |
| Masked Neighboring dMRI Correlation | `[raw|t1|t1post]_masked_neighbor_corr` | Higher value, closer to 1 |
| dMRI Contrast | `[raw|t1|t1post]_dwi_contrast` | Higher value |
| **Fiber orientation coherence** | | |
| Fiber Coherence Index | `[raw|t1|t1post]_coherence_index` | Higher value (better gradient validity) |
| Fiber Incoherence Index | `[raw|t1|t1post]_incoherence_index` | Lower value (better gradient validity) |
| **Slice artifact metrics** | | |
| Number of Bad Slices | `[raw|t1|t1post]_num_bad_slices` | Lower value (fewer slices with artifacts) |

**Supplementary Table S1 | Diffusion MRI quality metrics (IQMs) derived from *QSIPrep*.** Metrics summarize motion, signal quality, diffusion consistency, fiber orientation validity, and slice artifacts. Variable names correspond to *QSIPrep* outputs. Prefix notation [raw|t1|t1post] indicates the processing stage at which the metric was computed (raw diffusion data, preprocessed data prior to $B_1$ bias correction, or preprocessed data after $B_1$ bias correction).

| Model | Parameter | Description | Shells | Category |
|---|---|---|---|---|
| **qsirecon-DIPYDKI** | | | | |
| DKI | `ad` | Axial diffusivity | Full | Mean Diffusivity |
| DKI | `ak` | Axial kurtosis | Full | Diffusion Kurtosis |
| DKI | `fa` | Fractional anisotropy | Full | Diffusion Anisotropy |
| DKI | `kfa` | Kurtosis fractional anisotropy | Full | Diffusion Kurtosis |
| DKI | `md` | Mean diffusivity | Full | Mean Diffusivity |
| DKI | `mk` | Mean kurtosis | Full | Diffusion Kurtosis |
| DKI | `mkt` | Mean kurtosis tensor | Full | Diffusion Kurtosis |
| DKI | `rd` | Radial diffusivity | Full | Mean Diffusivity |
| DKI | `rk` | Radial kurtosis | Full | Diffusion Kurtosis |
| **qsirecon-DSIStudio** | | | | |
| GQI | `gfa` | Generalized fractional anisotropy | Full | Diffusion Anisotropy |
| GQI | `iso` | Isotropic diffusion | Full | |
| GQI | `qa` | Quantitative anisotropy | Full | Diffusion Anisotropy |
| Tensor | `ad` | Axial diffusivity (first eigenvalue) from a tensor fit | Inner | Mean Diffusivity |
| Tensor | `fa` | Fractional anisotropy from a tensor fit | Inner | Diffusion Anisotropy |
| Tensor | `ha*` | Helix angle from tensor fit | Inner | |
| Tensor | `md` | Mean diffusivity from a tensor fit | Inner | Mean Diffusivity |
| Tensor | `rd` | Radial diffusivity from a tensor fit | Inner | Mean Diffusivity |

| Model | Parameter | Description | Shells | Category |
|---|---|---|---|---|
| Tensor | rd1 | Lambda 2 (second eigenvalue) from a tensor fit | Inner | Mean Diffusivity |
| Tensor | rd2 | Lambda 3 (third eigenvalue) from a tensor fit | Inner | Mean Diffusivity |
| Tensor | txx* | Tensor fit txx | Inner | |
| Tensor | txy* | Tensor fit txy | Inner | |
| Tensor | txz* | Tensor fit txz | Inner | |
| Tensor | tyy* | Tensor fit tyy | Inner | |
| Tensor | tyz* | Tensor fit tyz | Inner | |
| Tensor | tzz* | Tensor fit tzz | Inner | |
| | | **qsirecon-TORTOISE_model-MAPMRI** | | |
| MAP-MRI | ng | Non-Gaussianity | Full | Diffusion Kurtosis |
| MAP-MRI | ngpar | Non-Gaussianity parallel | Full | Diffusion Kurtosis |
| MAP-MRI | ngperp | Non-Gaussianity perpendicular | Full | Diffusion Kurtosis |
| MAP-MRI | pa | Propagator anisotropy | Full | Diffusion Kurtosis |
| MAP-MRI | path | Propagator anisotropy (θ) | Full | Diffusion Kurtosis |
| MAP-MRI | rtap | Return to axis probability | Full | Mean Diffusivity |
| MAP-MRI | rtop | Return to origin probability | Full | Mean Diffusivity |
| MAP-MRI | rtpp | Return to plane probability | Full | Mean Diffusivity |
| Tensor | ad | Axial diffusivity | Inner | Mean Diffusivity |
| Tensor | am* | A0 from a tensor fit | Inner | |

| Model | Parameter | Description | Shells | Category |
|---|---|---|---|---|
| Tensor | `fa` | Fractional anisotropy from a tensor fit | Inner | Diffusion Anisotropy |
| Tensor | `li` | Lattice index | Inner | Diffusion Anisotropy |
| Tensor | `rd` | Radial diffusivity from a tensor fit | Inner | Mean Diffusivity |
| | | **qsirecon-wmNODDI** | | |
| NODDI | `icvf` | Intracellular volume fraction | Full | Diffusion Kurtosis |
| NODDI | `isovf` | Isotropic volume fraction | Full | Mean Diffusivity |
| NODDI | `od` | Orientation dispersion index | Full | Diffusion Anisotropy |

**Supplementary Table S2 | Diffusion MRI microstructural metrics derived from *QSIRecon* reconstruction pipelines.** Diffusion MRI metrics generated from multiple *QSIRecon* reconstruction workflows, including *DIPY* diffusion kurtosis imaging (DKI), *DSI Studio* generalized *q*-sampling imaging (GQI), *TORTOISE* mean apparent propagator MRI (MAP-MRI), tensor fitting, and neurite orientation dispersion and density imaging (NODDI). The *Model* column indicates the reconstruction framework used to estimate each metric, while the *Parameter* column lists the corresponding output variable name from the reconstruction pipeline. The *Description* column provides a brief definition of each metric, and the *Shells* column indicates whether the metric was estimated using all diffusion shells (*Full*) or only the inner shell (*b < 1,250*; *Inner*) of the dMRI acquisition. Where applicable, the categorization from Sadikov et al., (2025)[1] is included as a guideline for general interpretation of each parameter (each Category is shaded with a different color). There were a total of 14 “Mean Diffusivity” (beige) measures, 7 “Diffusion Anisotropy” (green) measures, 11 “Diffusion Kurtosis” (purple) measures, and 9 uncategorized measures (gray). Parameters marked with an asterisk (*) were generated by the reconstruction pipeline but were not included in any downstream analyses. Readers are referred to Table 1 of Yeh, 2020[2] for definitions of macrostructural measures.

| Site | Scanner Vendor | Model | Device Serial Number (anonymized) | Major Software Version | Number of Scans |
|---|---|---|---|---|---|
| 1 | Philips | Achieva dStream | anonb2d4 | 5.3 | 320 |
| 1 | Philips | Achieva dStream | anonb2d4 | 5.4 | 154 |
| 1 | Philips | Achieva dStream | anonb2d4 | 5.7 | 120 |
| 2 | Siemens | Prisma Fit | anon2565 | VE11B | 484 |
| 2 | Siemens | Prisma Fit | anon2565 | VE11C | 353 |
| 2 | Siemens | Prisma Fit | anon2565 | VE11E | 537 |
| 3 | Siemens | Prisma | anon8928 | VE11C | 1251 |
| 4 | GE | Discovery MR750 | anon239d | DV25 | 229 |
| 4 | GE | Discovery MR750 | anon239d | DV26 | 447 |
| 4 | GE | Discovery MR750 | anonf61b | DV25 | 367 |
| 4 | GE | Discovery MR750 | anonf61b | DV26 | 705 |
| 5 | Siemens | Prisma Fit | anon02a4 | VE11C | 864 |
| 6 | Siemens | Prisma Fit | anonfaf6 | VE11B | 474 |
| 6 | Siemens | Prisma Fit | anonfaf6 | VE11C | 650 |
| 7 | Siemens | Prisma Fit | anon241f | VE11C | 550 |
| 8 | GE | Discovery MR750 | anon12da | DV25 | 88 |
| 8 | GE | Discovery MR750 | anon12da | DV26 | 422 |
| 8 | GE | Discovery MR750 | anon12da | DV29 | 75 |
| 9 | Siemens | Prisma Fit | anonc805 | VE11B | 16 |
| 9 | Siemens | Prisma Fit | anonc805 | VE11C | 724 |

| | | | | | |
|---|---|---|---|---|---|
| 10 | GE | Discovery MR750 | anon3255 | DV25 | 387 |
| 10 | GE | Discovery MR750 | anon3255 | DV26 | 596 |
| 10 | GE | Discovery MR750 | anon3255 | DV29 | 260 |
| 10 | GE | Discovery MR750 | anon9d57 | DV26 | 406 |
| 11 | Siemens | Prisma | anondc7b | VE11C | 852 |
| 12 | Siemens | Prisma Fit | anon6ecd | VE11C | 45 |
| 12 | Siemens | Prisma Fit | anonefd2 | VE11C | 1053 |
| 12 | Siemens | Prisma Fit | anonefd2 | VE11E | 250 |
| 13 | GE | Discovery MR750 | anon2b69 | DV25 | 103 |
| 13 | GE | Discovery MR750 | anon2b69 | DV26 | 163 |
| 13 | GE | Discovery MR750 | anonbcc2 | DV25 | 387 |
| 13 | GE | Discovery MR750 | anonbcc2 | DV26 | 303 |
| 13 | GE | Discovery MR750 | anoncccd | DV26 | 187 |
| 13 | GE | Discovery MR750 | anoncccd | DV29 | 76 |
| 13 | GE | SIGNA UHP | anonbcc2 | RX28 | 345 |
| 14 | Siemens | Prisma | anon3eba | VE11C | 1221 |
| 14 | Siemens | Prisma Fit | anona429 | VE11B | 17 |
| 14 | Siemens | Prisma Fit | anona429 | VE11C | 195 |
| 15 | Siemens | Prisma Fit | anon7a3b | VE11B | 292 |
| 15 | Siemens | Prisma Fit | anon7a3b | VE11C | 498 |
| 16 | Siemens | Prisma | anon0bd7 | VE11B | 717 |
| 16 | Siemens | Prisma | anon0bd7 | VE11C | 1578 |

| | | | | | |
|---|---|---|---|---|---|
| 17 | Philips | Achieva dStream | anond450 | 5.3 | 301 |
| 17 | Philips | Achieva dStream | anond450 | 5.4 | 253 |
| 17 | Philips | Achieva dStream | anond450 | 5.6 | 158 |
| 17 | Philips | Achieva dStream | anond450 | 5.7 | 200 |
| 18 | GE | Discovery MR750 | anon0982 | DV25 | 86 |
| 18 | GE | Discovery MR750 | anon0982 | DV26 | 319 |
| 18 | GE | Discovery MR750 | anon6d38 | DV26 | 120 |
| 18 | GE | SIGNA Premier | anone411 | RX28 | 328 |
| 19 | Philips | Ingenia | anon1364 | 5.3 | 442 |
| 19 | Philips | Ingenia | anon1364 | 5.6 | 114 |
| 19 | Philips | Ingenia | anon1364 | 5.7 | 216 |
| 20 | Siemens | Prisma | anon224d | VE11C | 1392 |
| 20 | Siemens | Prisma | anond7c2 | VE11C | 15 |
| 20 | Siemens | Prisma Fit | anon7353 | VE11B | 17 |
| 20 | Siemens | Prisma Fit | anon7353 | VE11C | 161 |
| 21 | Siemens | Prisma | anon1f4f | VE11C | 608 |
| 21 | Siemens | Prisma | anon91db | VE11C | 76 |
| 21 | Siemens | Prisma Fit | anonbe03 | VE11B | 611 |

**Supplementary Table S3 | Scanner information for harmonization batches.** Each batch corresponds to a unique combination of device serial number and major software version. The number of scans per batch is reported. Batches of fewer than 15 scans were excluded from analysis and not included in this table.

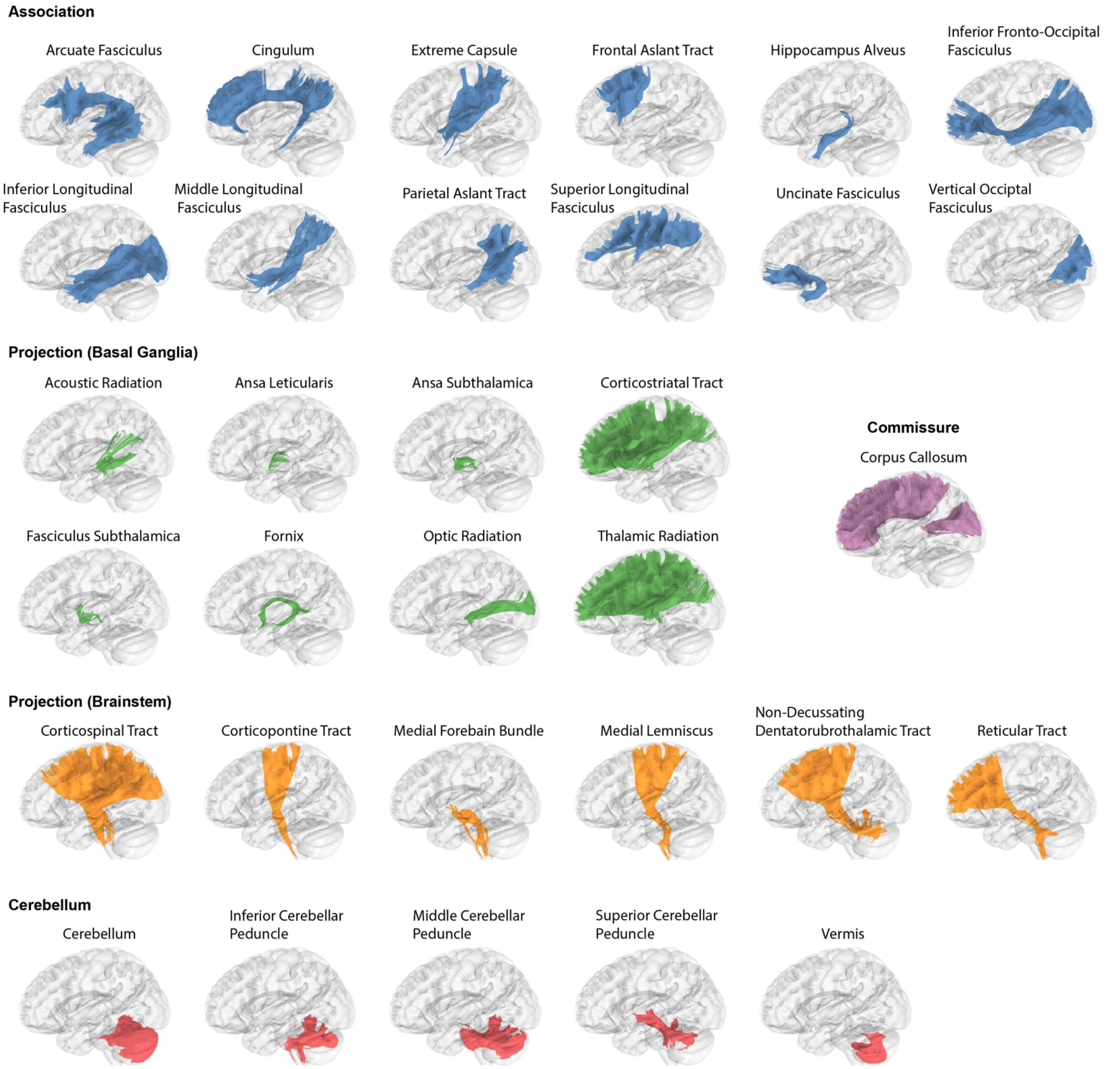


**Supplementary Figure S1 | White matter bundles reconstructed using *DSI Studio AutoTrack*.** Template bundle reconstructions of white matter tracts generated using *DSI Studio AutoTrack,* grouped by anatomical class. For bilateral bundles, only the left hemispheric homolog is shown.

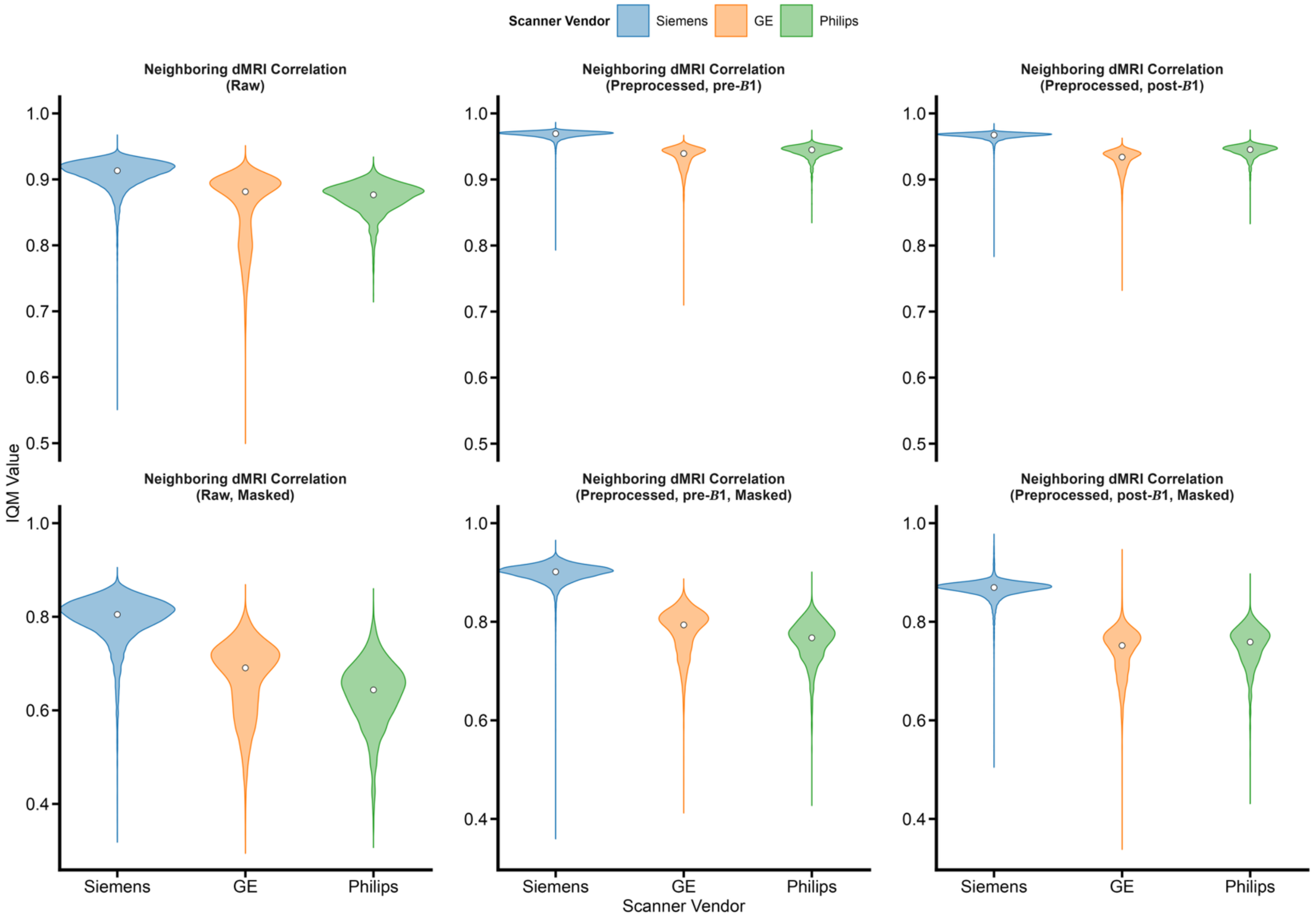


**Supplementary Figure S2 | Neighboring dMRI correlation across scanner vendors.** Violin plots are colored by scanner vendor. White dots represent the vendor median. All vendor pairwise comparisons were significant at $q < 0.05$.

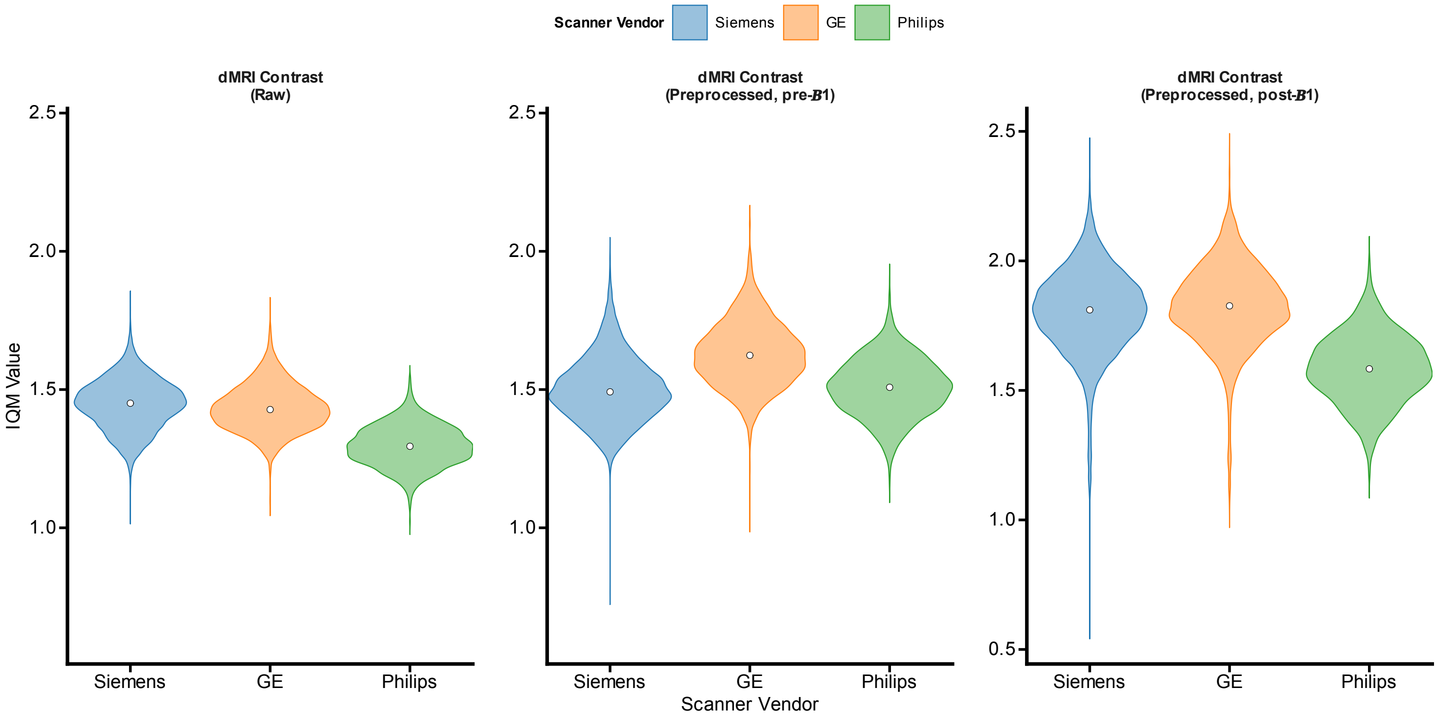


**Supplementary Figure S3 | dMRI contrast values across scanner vendors.** Violin plots are colored by scanner vendor. White dots represent the vendor median. All vendor pairwise comparisons were significant at $q < 0.05$.

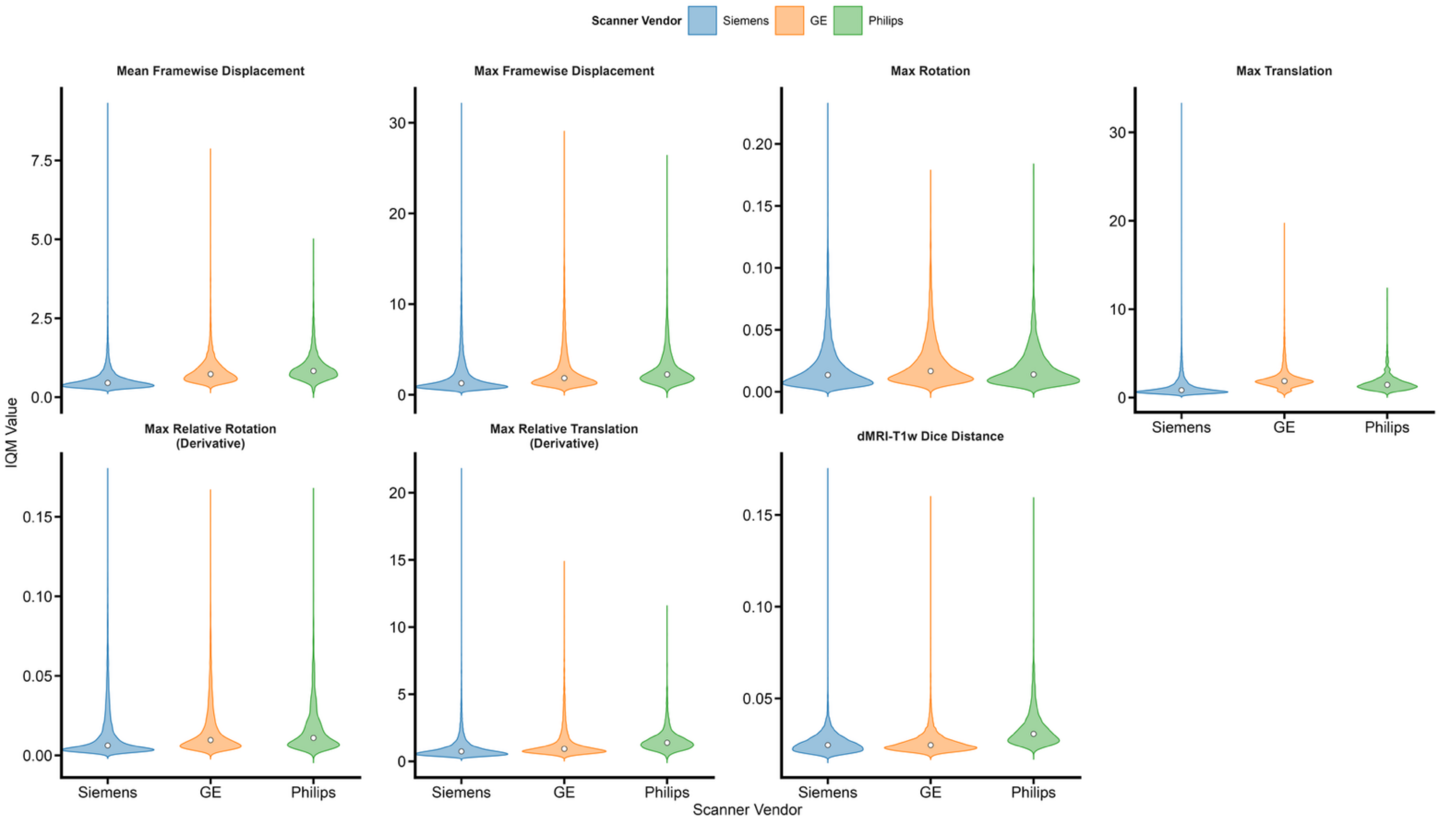


**Supplementary Figure S4 | Motion and co-registration across scanner vendors.** Violin plots are colored by scanner vendor. White dots represent the vendor median. All vendor pairwise comparisons were significant at $q < 0.05$.

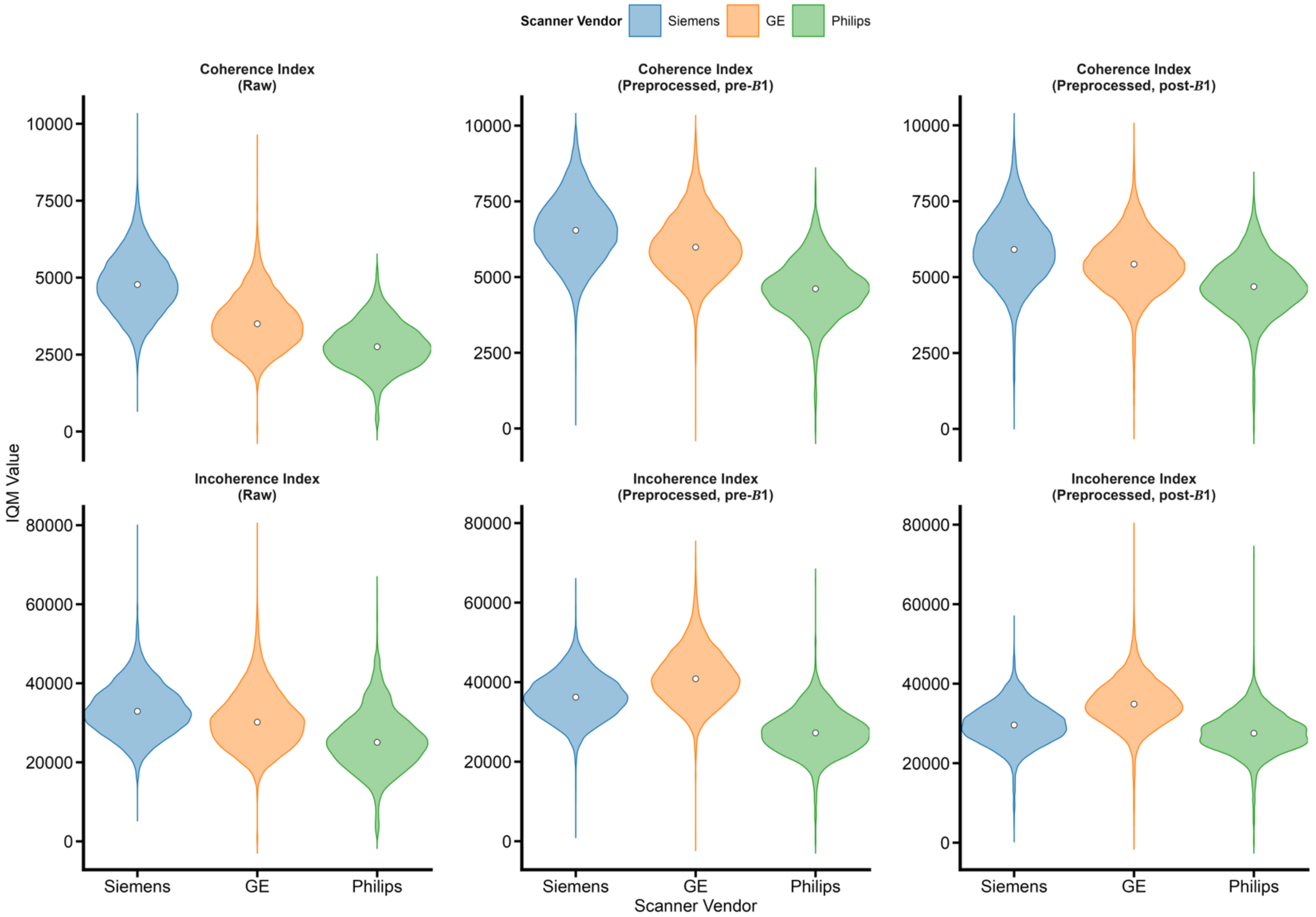


**Supplementary Figure S5 | Fiber incoherence and incoherence indices across scanner vendors.** Violin plots are colored by scanner vendor. White dots represent the vendor median. All vendor pairwise comparisons were significant at $q < 0.05$.

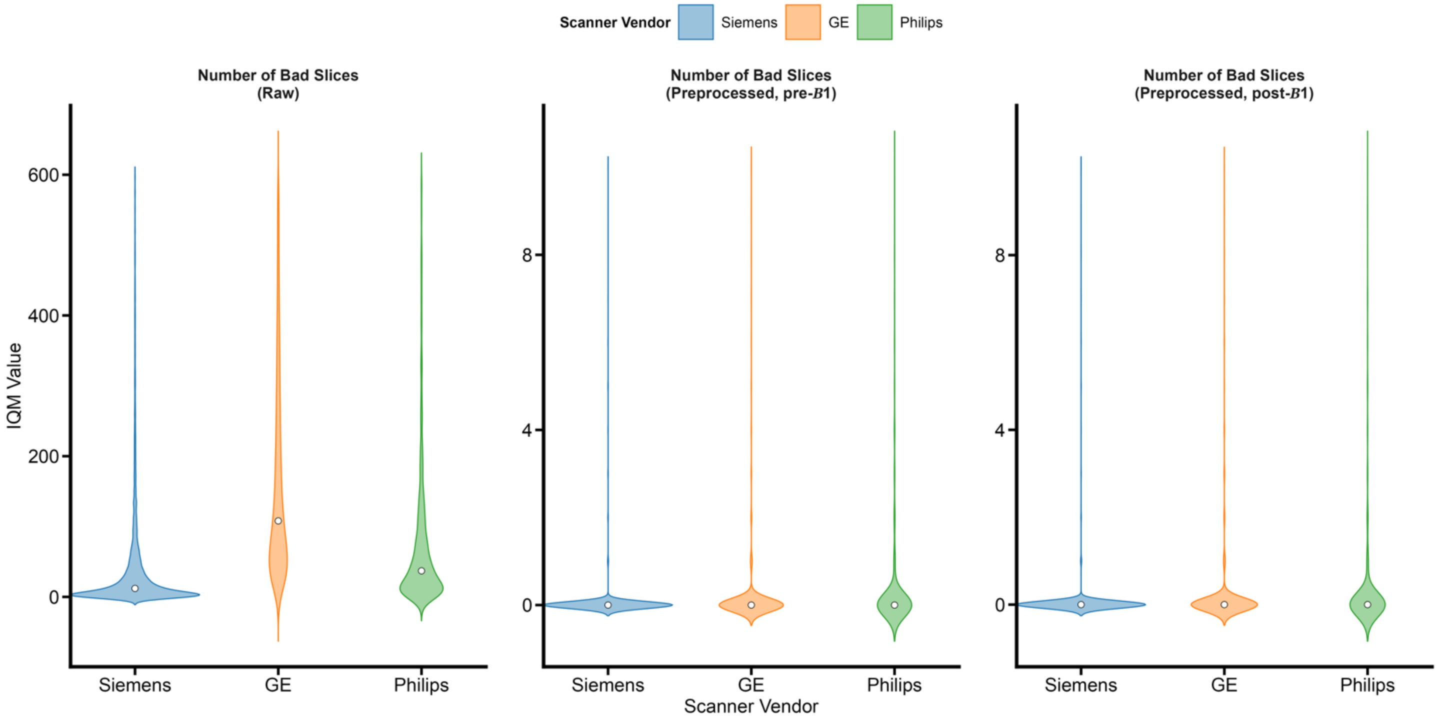


**Supplementary Figure S6 | Number of bad dMRI slices across scanner vendors.** Violin plots are colored by scanner vendor. White dots represent the vendor median. All vendor pairwise comparisons were significant at $q < 0.05$.

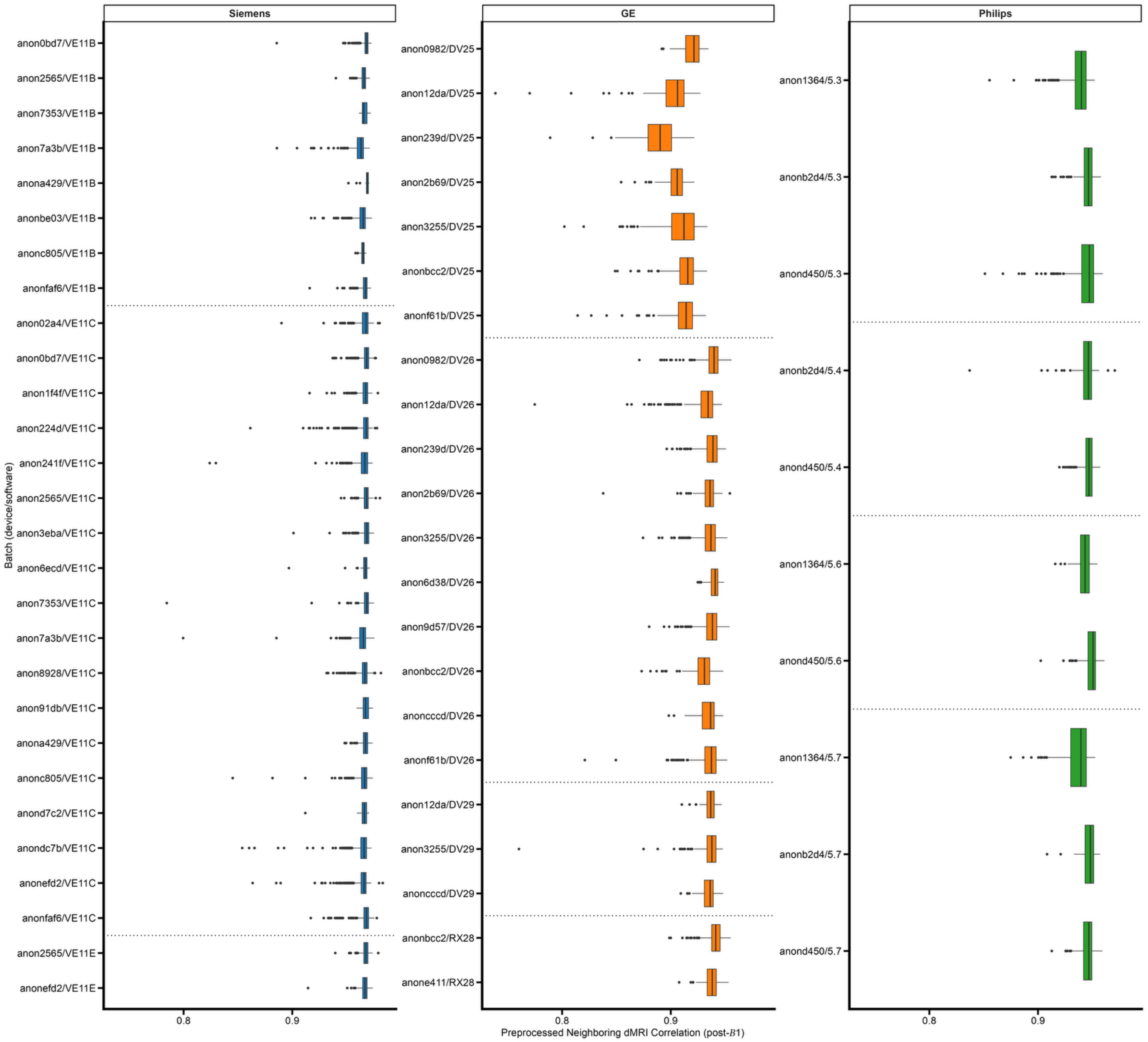


**Supplementary Figure S7 | Neighboring dMRI correlation across batches.** Each batch is a unique device-software version combination. Box plots are colored by vendor. Vertical black lines denote the median value. Dotted horizontal lines separate software versions.

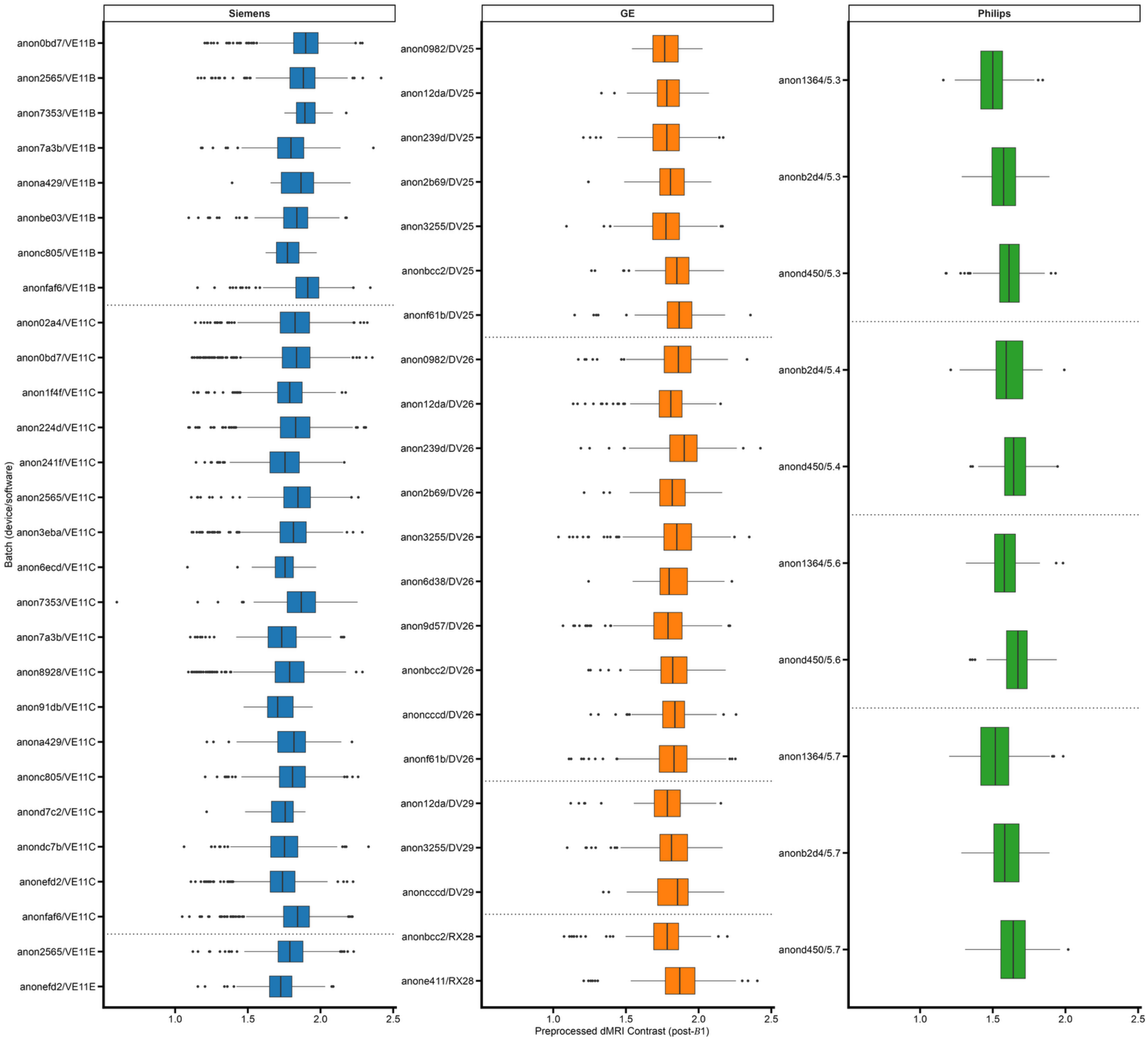


**Supplementary Figure S8 | dMRI contrast values across batches.** Each batch is a unique device-software version combination. Box plots are colored by vendor. Vertical black lines denote the median value. Dotted horizontal lines separate software versions.

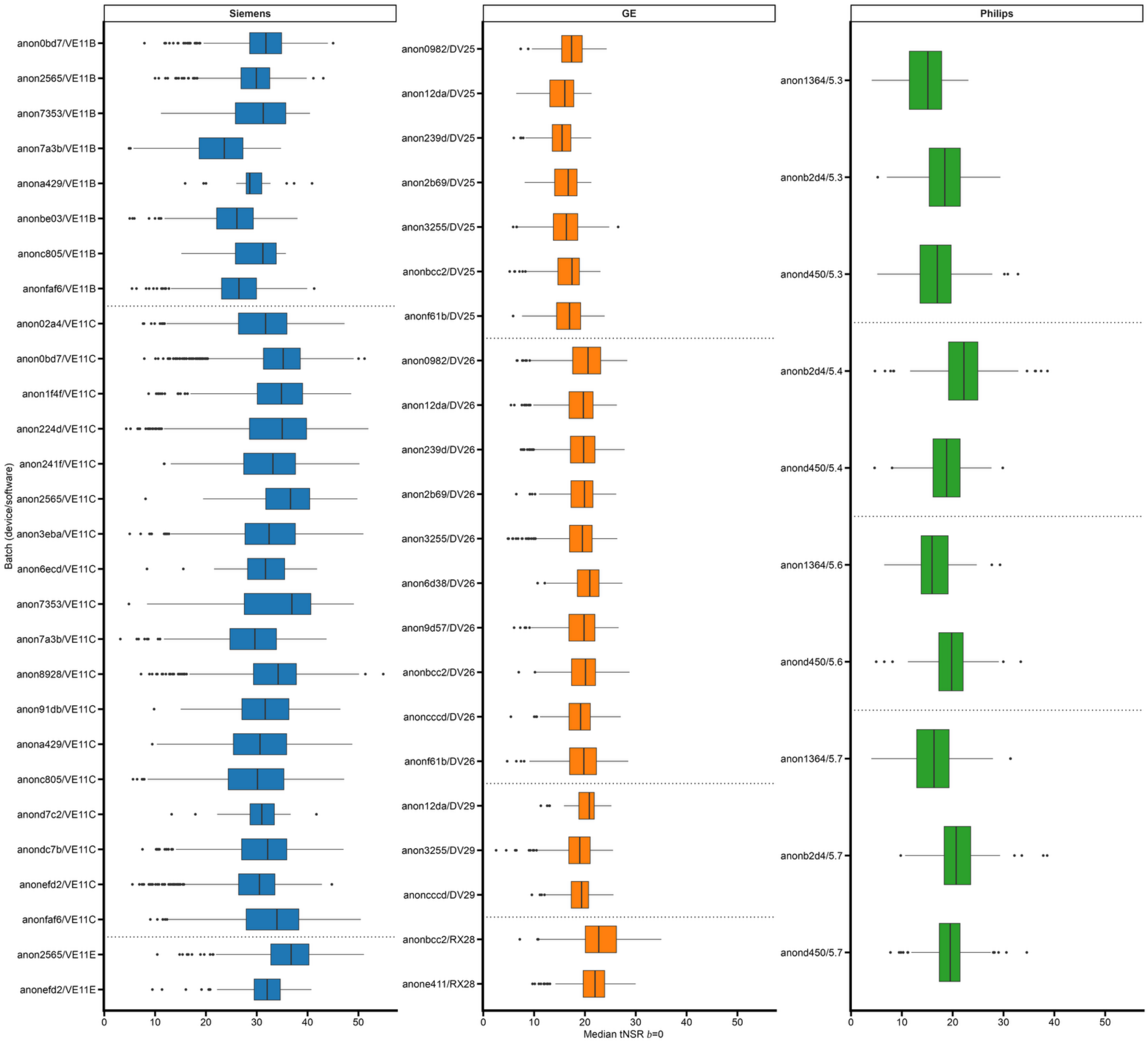


**Supplementary Figure S9 | median *b* = 0 tSNR values across batches.** Each batch is a unique device-software version combination. Box plots are colored by vendor. Vertical black lines denote the median value. Dashed horizontal lines separate software versions.

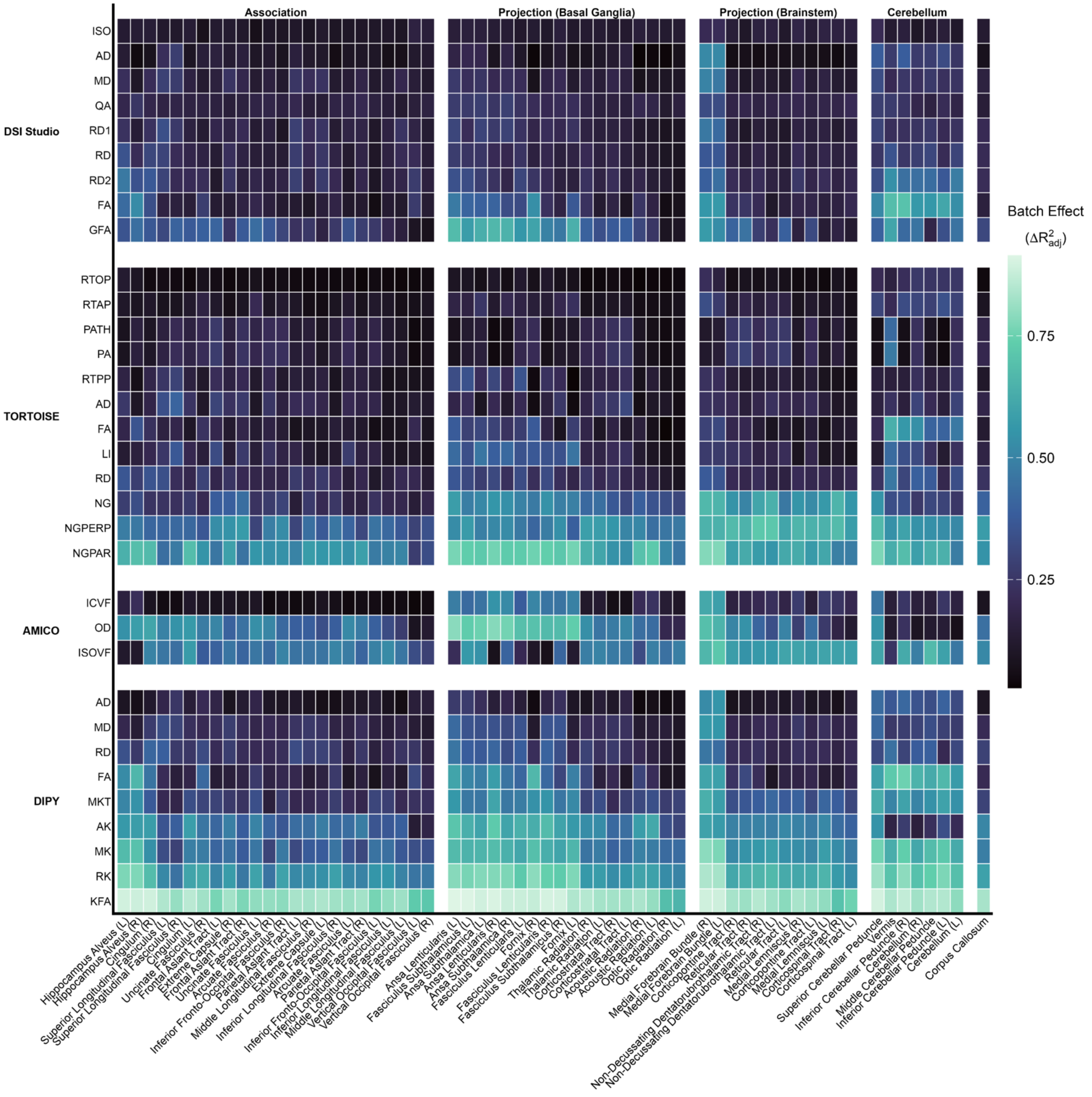


**Supplementary Figure S10 | Batch effects across microstructural metrics and white matter bundles.** Columns are grouped by bundle category, and rows are grouped by processing software. Within each block, rows and columns are ordered by average batch effect size.

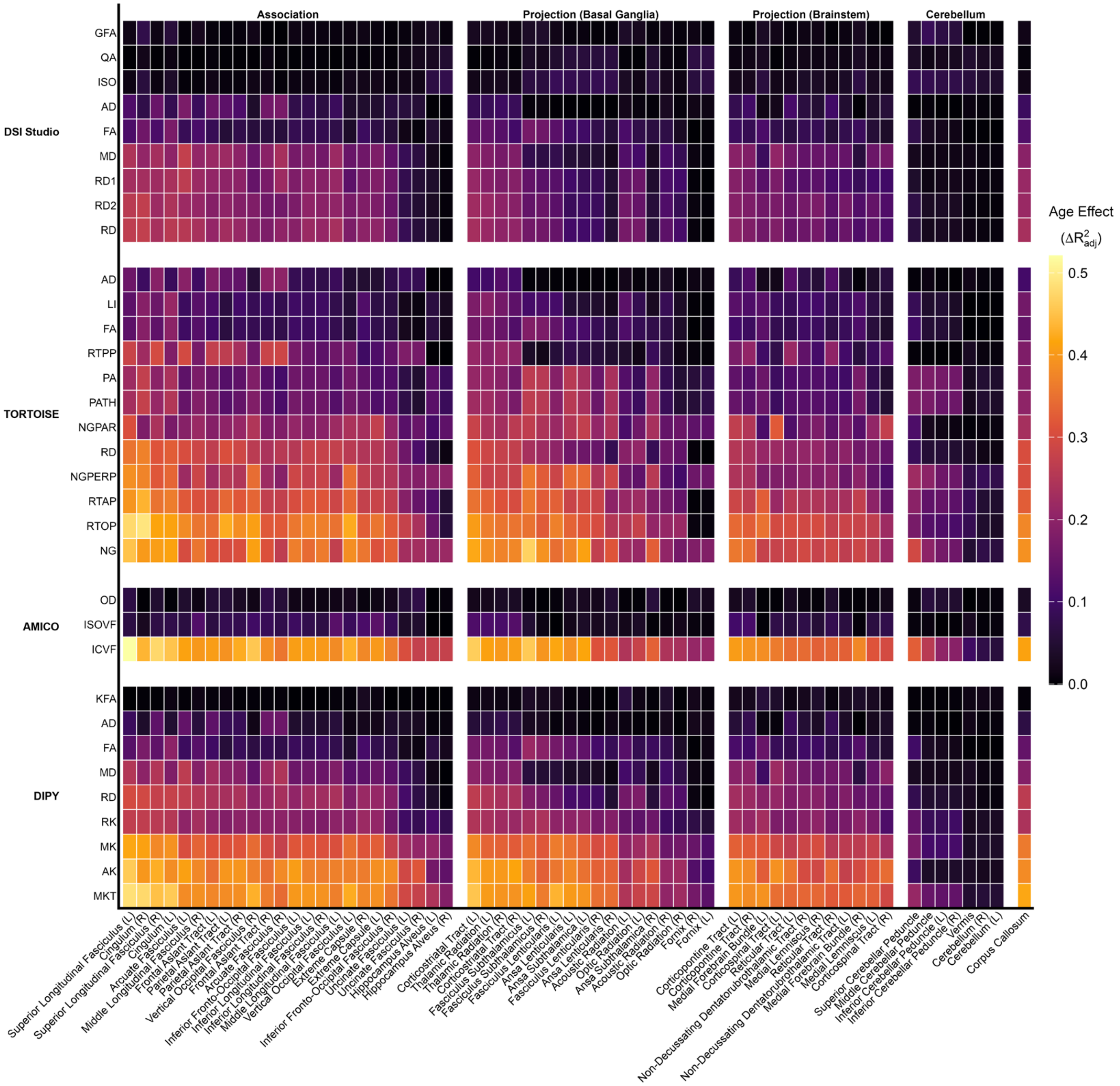


**Supplementary Figure S11 | Age effects across microstructural metrics and white matter bundles.** Columns are grouped by bundle category, and rows are grouped by processing software. Within each block, rows and columns are ordered by average age effect size.

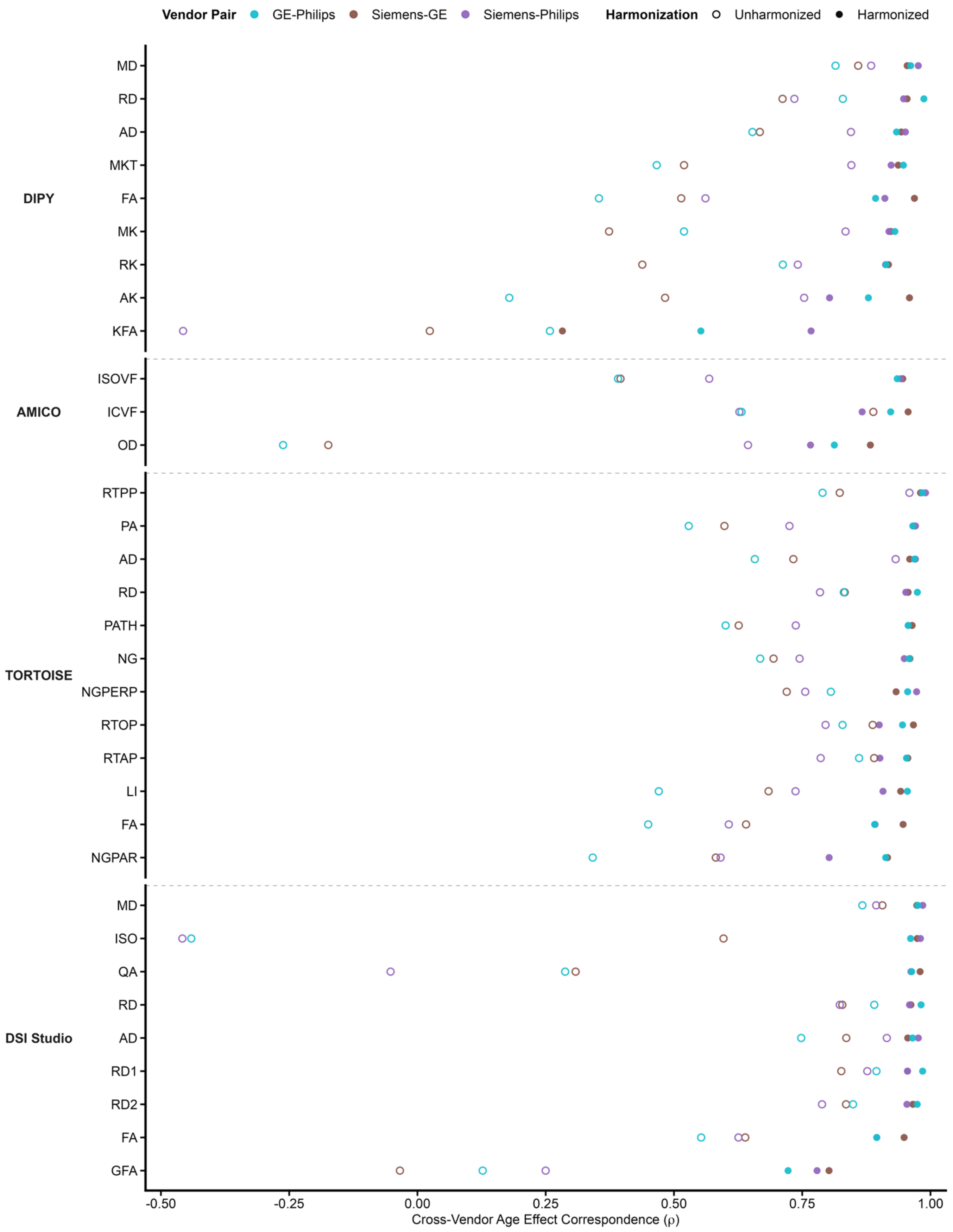


**Supplementary Figure S12 | Cross-vendor age effect correspondence across all white matter metrics.** Microstructural metrics are grouped by software, separated by grey dashed lines. Within software, metrics are ordered by average harmonized cross-vendor correspondence across the 3 pairwise comparisons. Dots are colored by vendor pair, with different fill patterns for unharmonized and harmonized versions of the metric.

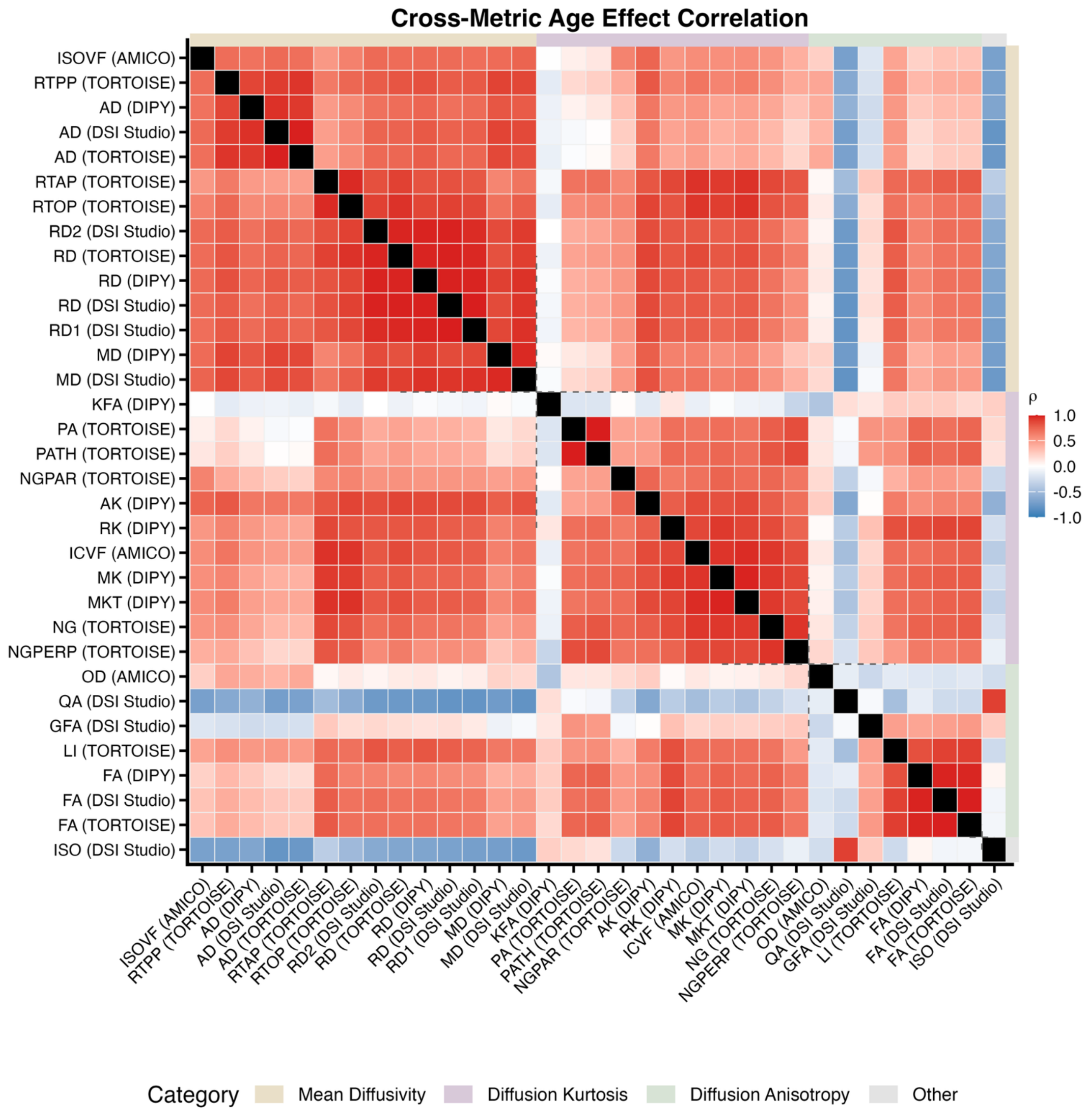


**Supplementary Figure S13 | Cross-metric age effect correspondences across all white matter metrics.** Pairwise Spearman correlations are shown between bundle-wise age-effect vectors for all microstructural metrics. Metrics are grouped by microstructural category (defined in **Supplementary Table S2**; beige - Mean Diffusivity, purple - Diffusion Kurtosis, green - Diffusion Anisotropy), with category blocks indicated by colored axis strips; within each category, metrics are ordered by hierarchical clustering of age-effect profiles (average linkage; distance = 1 − |ρ|) so metrics with similar developmental patterns are adjacent.

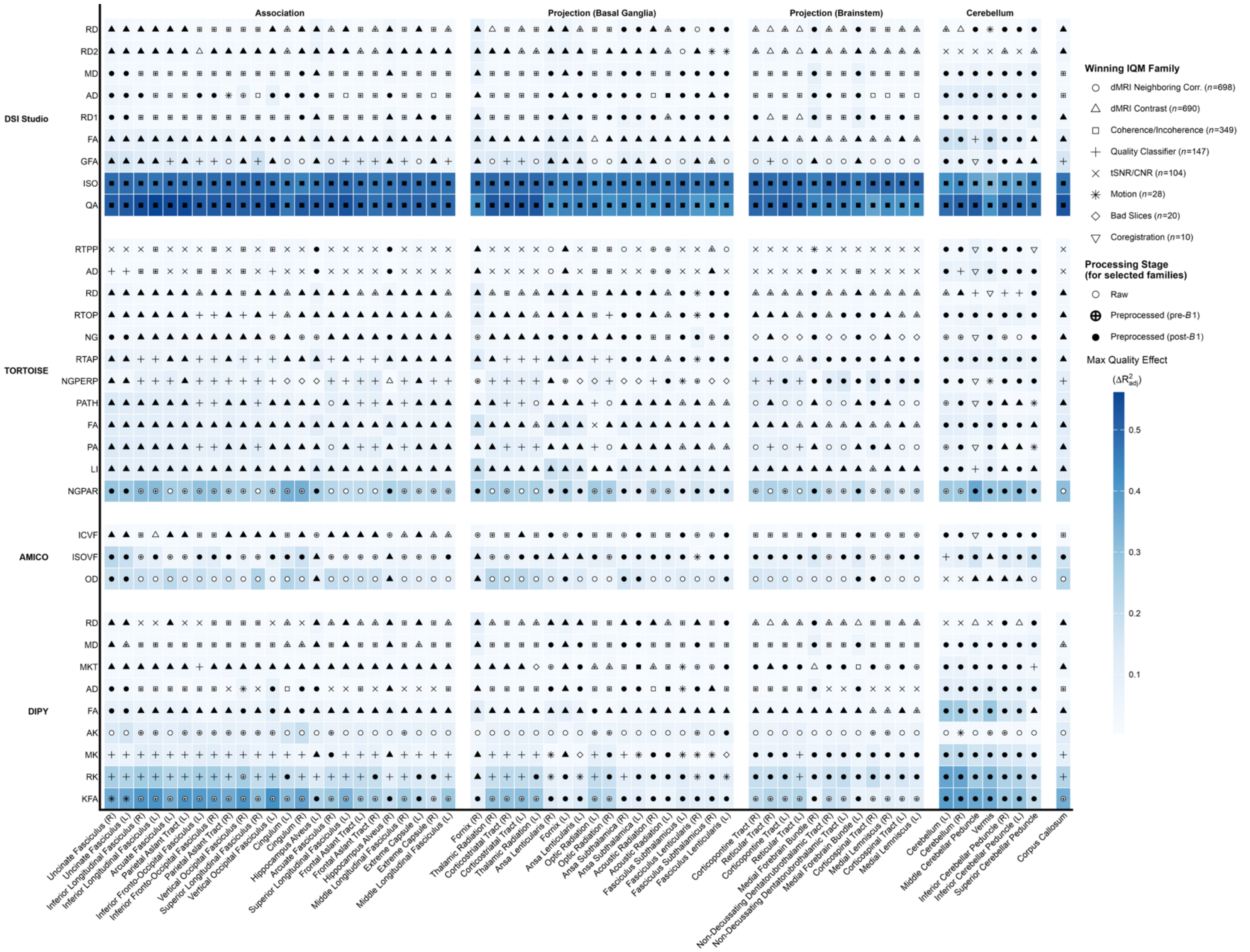


**Supplementary Figure S14 | Quality effects across microstructural metrics and white matter bundles.** Each cell is colored by the highest quality effect achieved in that bundle–microstructure combination. Cell symbols denote which quality metric family the highest effect metric belonged to. When relevant, marker fillings denote which processing stage the quality metric was derived from. Columns are grouped by bundle category, and rows are grouped by processing software. Within each block, rows and columns are ordered by average max quality effect size.

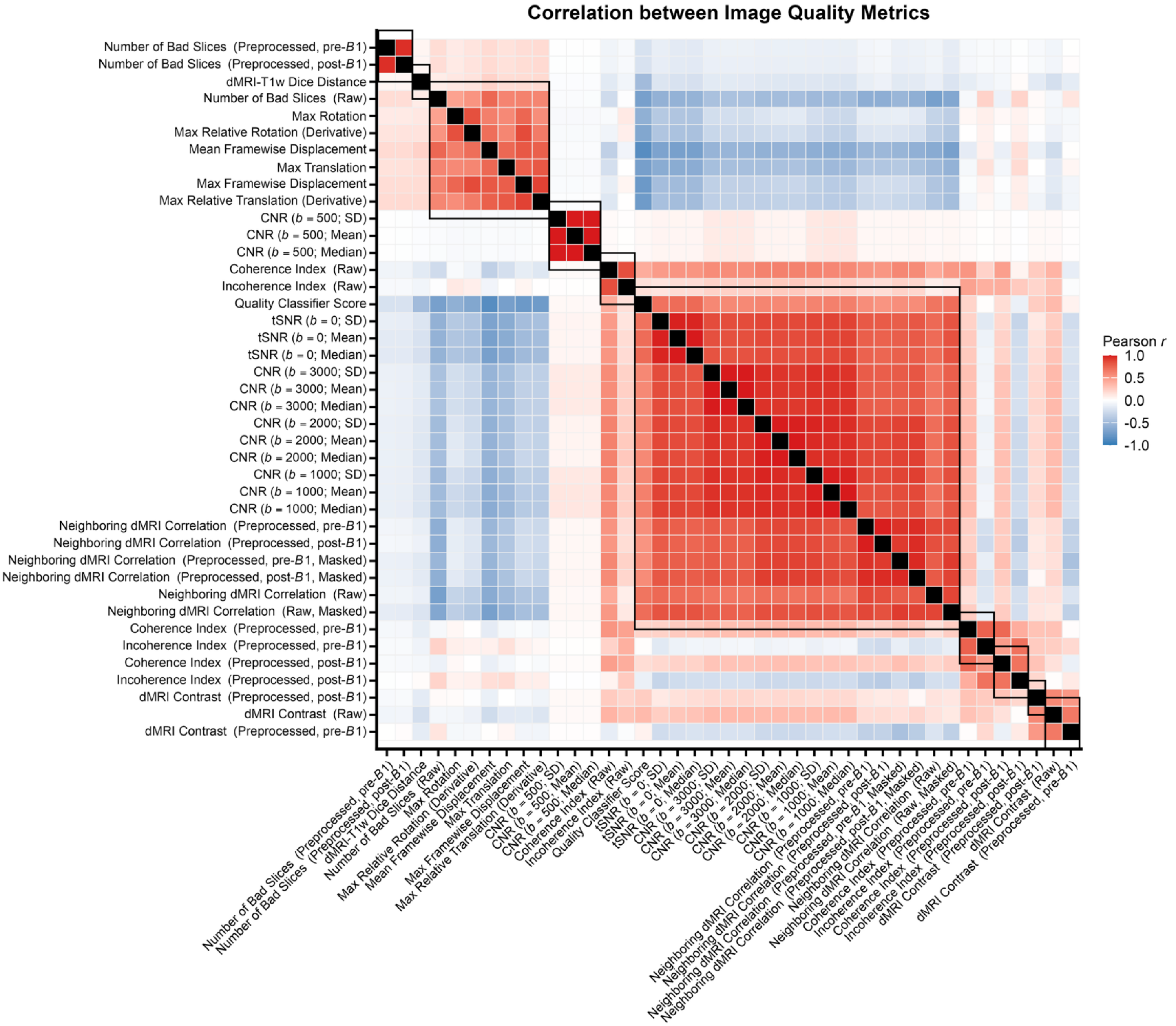


**Supplementary Figure S15 | Correlation between image quality metrics.** Each cell shows the pairwise Pearson correlation between IQMs across sessions. IQMs are grouped by quality family, with IQM family blocks ordered from largest to smallest; within each family, IQMs are ordered by hierarchical clustering of their correlation structure (average linkage; distance = 1 − *r*), so covarying IQMs are adjacent.

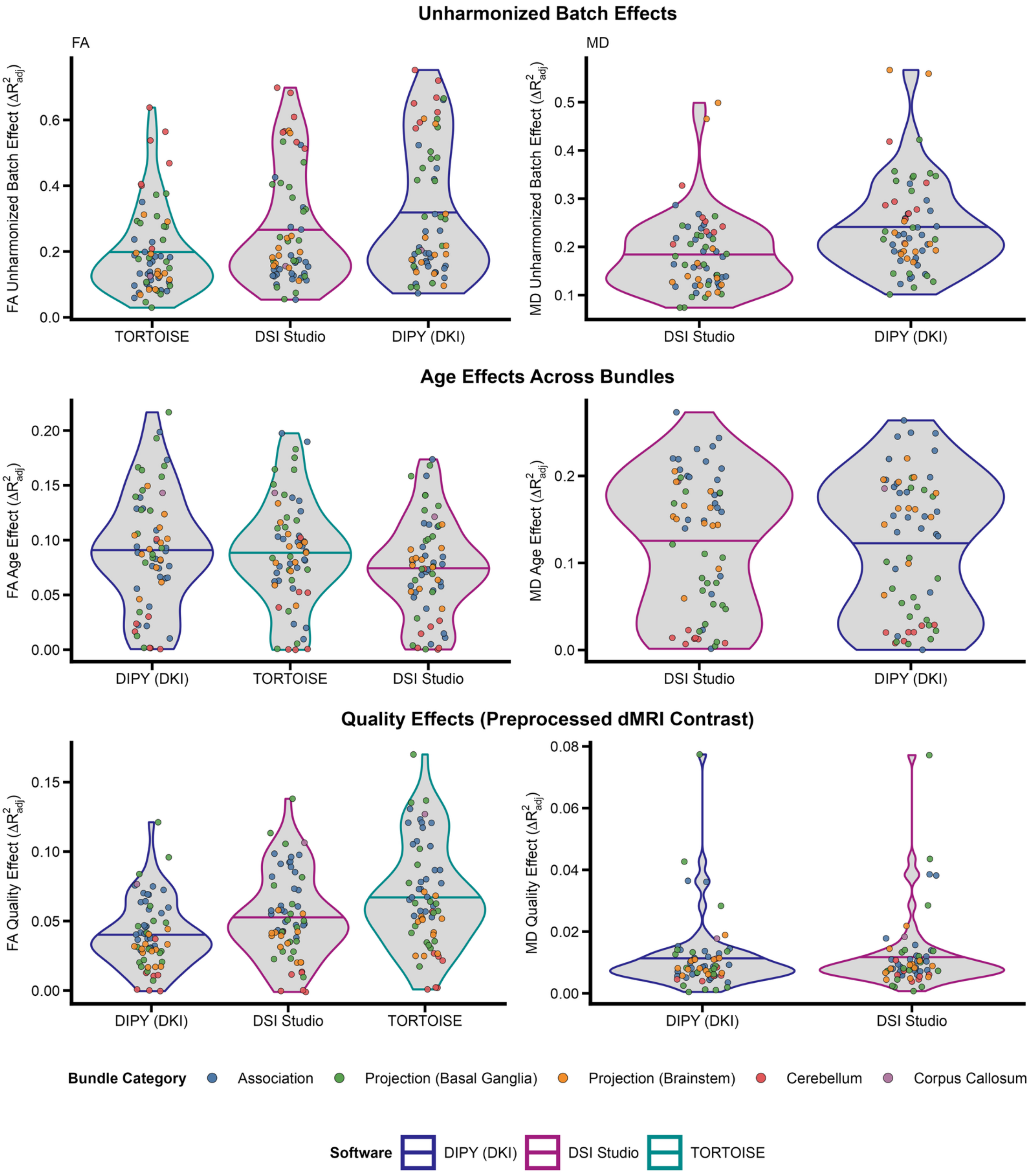


**Supplementary Figure S16 | Comparison of batch, age, and quality effects across tensor-fitting workflows.** Violins are colored by tensor-fitting software. Violins are ordered by the average effect size, denoted by the horizontal line in each violin. Each dot is a bundle, colored by its bundle category. The left column focuses on fractional anisotropy (FA) and the right column focuses on mean diffusivity (MD). The MAP-MRI model did not produce a mean diffusivity measure.

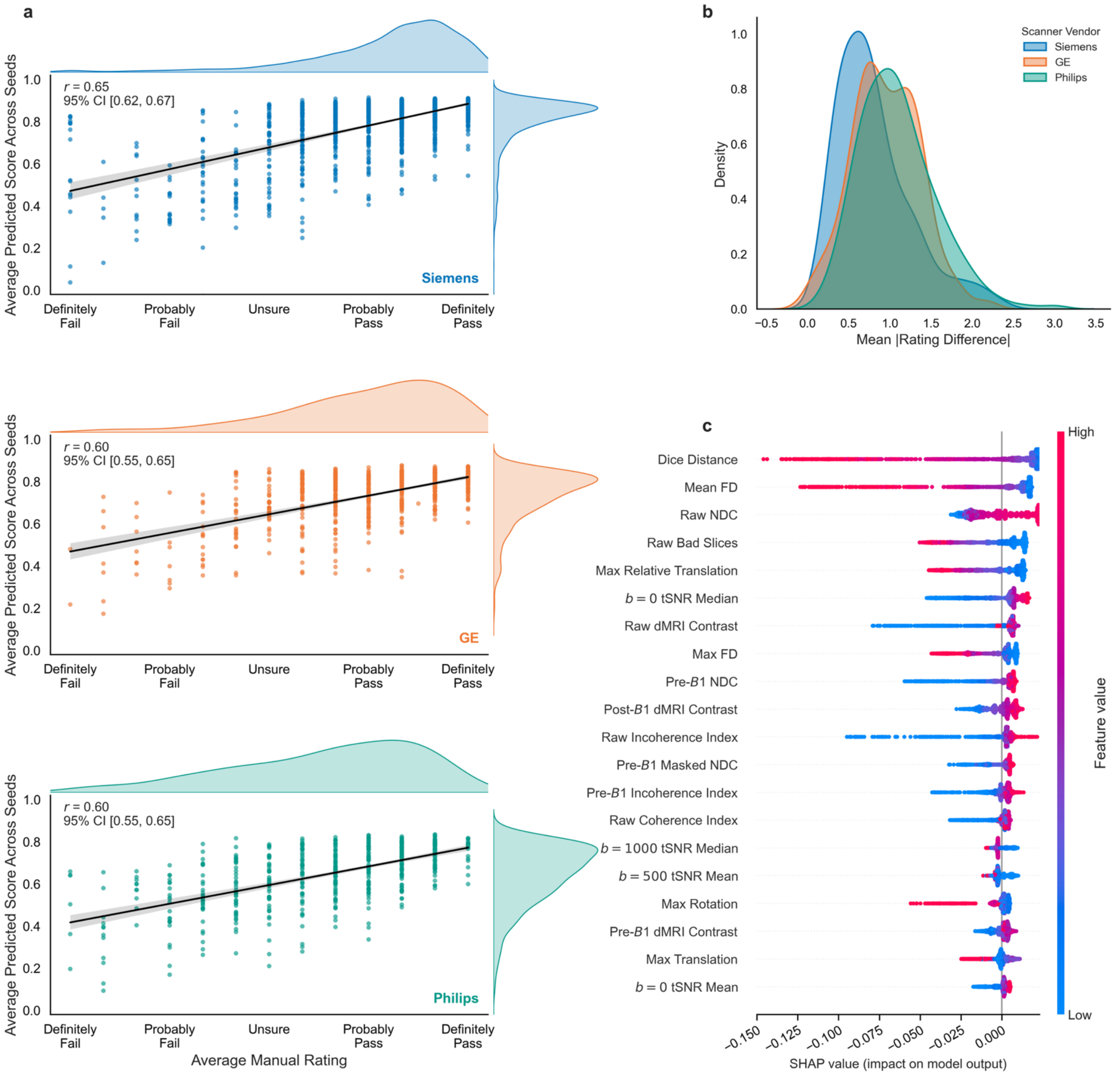


**Supplementary Figure S17 | Quality ratings, classification, rater disagreement, feature importance. (a)** Mean manual image-quality rating plotted against mean model-predicted quality score (averaged across 1,000 cross-validation seeds), with black linear fit lines and 95% confidence intervals; marginal density plots are shown for both axes. **(b)** Inter-rater disagreement distributions by scanner vendor. Disagreement for each image and pair of reviewers is quantified as the absolute difference in rating scores between reviewer pairs. **(c)** SHAP beeswarm summary for the final quality-classification model, showing feature-level contributions to the model. Only the top 20 features are shown here.

#### Supplementary Note S1: Automated quality classification

We trained a regression-based machine-learning model to predict manual dMRI quality ratings from the 40 image quality metrics (IQMs) produced by *QSIPrep*. For each dMRI image, the means of the three independent manual ratings were used as the prediction targets. Following prior work on automated dMRI quality assessment by Richie-Halford, Cieslak, and colleagues[3], we used an eXtreme Gradient Boosting (XGBoost) model[4]; however, instead of predicting a binary pass/fail label, we implemented an XGBRegressor classifier to predict the continuous mean quality rating. This approach preserved the variability in image quality and yielded a continuous prediction metric that we used directly as a covariate in subsequent statistical analyses. Images from all scanner vendors were pooled for model training.

To evaluate model performance and mitigate overfitting, we used a nested 5×5-fold cross-validation framework. Both inner and outer folds were stratified to balance for scanner-vendor representation and the proportion of ratings above and below 0. Data in the inner folds were used to tune hyperparameters, which were optimized using a 5-fold cross validation Bayesian search procedure (200 iterations). The model configuration yielding the lowest mean cross-validated root mean squared error (RMSE) across the inner folds was selected as the best-performing model and then evaluated on the held-out outer test fold. This entire process was repeated across 1,000 random seeds, yielding 1,000 predicted quality scores per rated image. To quantify the performance of the model, we ran a Pearson correlation between the average of the 3 manual ratings and the average of the 1,000 predicted quality scores (**Supplementary Fig. S17a**).

After cross-validation performance was established, a final model was retrained on all labeled data (i.e., images with manual quality ratings). Hyperparameters were similarly optimized using a Bayesian search with 200 iterations, and the best-performing model was retained. This final trained model was then applied to all unlabeled images in the dataset to generate predicted quality scores based on their *QSIPrep*-derived IQMs. To prevent overfitting, each labeled image was assigned a quality score equal to the average of the 1,000 predictions during cross-validation. Feature importance for each IQM was assessed using SHapley Additive exPlanations (SHAP) values (**Supplementary Fig. S17c**)

#### Supplementary Note S2: Tract geometry plausibility check

To assess the reliability and anatomical plausibility of tractography generated by *DSI Studio*'s *AutoTrack*, we conducted a series of sanity checks based on expected geometric properties of bundle reconstructions. We focused on four white matter bundles that are consistently identifiable and informative about reconstruction quality: the corpus callosum (CC), bilateral arcuate fasciculus (AF), bilateral corticospinal tract (CST), and bilateral inferior fronto-occipital fasciculus (IFOF). For each dataset, we first confirmed the presence of each bundle. Datasets in which any of these bundles were incomplete or had a total volume below 100 $mm^3$ were flagged, as such values are implausibly small. We then examined the following tract geometry metrics[2] that describe shape and consistency across hemispheres. Streamline curl reflects how much a tract bends or twists. The length ratio measures the symmetry of bundle length between left and right hemispheres; values near 1 indicate good symmetry. The volume ratio compares the total spatial extent of left and right bundles, providing a coarse measure of reconstruction balance. Based on anatomical priors, extensive review of individual-level data, and empirical distributions, we defined very conservative heuristics to identify suspect data. The CC should have more curl than the CST and IFOF, which are both relatively straight bundles, i .e.,: curl(CC–CST) > 0 and curl(CC–IFOF) > 0. We also would expect bilateral bundle homologs to be nearly symmetrical, i.e., AF length ratio < 2, CST and IFOF length ratios < 1.25, and volume ratios < 5. Datasets failing any of these criteria were marked for exclusion from all downstream analyses. In total, 310 datasets were flagged for exclusion by this process.
